# Differentiation and maturation of iPSC-derived motor and sensory neurospheres using biomodified PEG-based microgels

**DOI:** 10.64898/2026.08.19.745683

**Authors:** Laura Klasen, Céline Bastard, Matthias Mork, Greta Romahn, José Luis Gerardo-Nava, Laura De Laporte

## Abstract

Sensory and motor neurons differ significantly in their morphology, structural organization, functional properties, and mode of action. However, despite this heterogeneity, many *in vitro* studies focus only on a single neuronal subtype, mainly being sensory neurons, limiting the translational potential and relevance of these studies for the evaluation of therapeutic options for spinal cord injury. In this study, we investigate the differentiation, maturation, and neuronal outgrowth of human induced pluripotent stem cell (iPSC)-derived motor and sensory neurospheres using polyethylene glycol (PEG)-microgels with various bioactive coatings. Our results show subtype-specific responses to the PEG-microgel scaffolds, with respect to motor and sensory neurosphere morphology and size. Furthermore, we compare the formation of the PEG-microgel/scaffolds when starting from iPSCs-derived precursor neuron spheres versus undifferentiated iPSCs. We observe notable differences in structural organization, maturity, and neuronal outgrowth between the two approaches, as well as between motor and sensory neurospheres. Together, these results underline the importance of studying motor and sensory neurons separately and highlight the need for a controlled, tunable culture platform to assess the impact of the microenvironment and to improve the physiological relevance of *in vitro* platforms for neuron-based research.

## 1. Introduction

Although motor and sensory neurons are essential to neural information processing, comparative studies of their structural, functional, and developmental properties remain limited. To enable effective regeneration after SCI, both sensory neuron input and motor neuron output must be restored. Therefore, the sensory neurons must re-establish ascending connections across the lesion site, while the descending motor neuron pathways, with their origin rostral to the injury, must reconnect with the spinal motor circuits. Supporting sensory and motor neuron regeneration is essential to reconnect the required sensorimotor circuits with their native targets.^[1,2]^ Motor and sensory neurons vary in functional polarity, anatomical localization, axonal architecture, and regenerative capacity, distinctions that are highly relevant to the design of biomaterials aimed at nerve repair, guidance, and functional integration.^[3,4]^

While motor neurons transmit signals from the central nervous system to peripheral tissues and possess cell bodies within the spinal cord or brainstem, sensory neurons convey information from peripheral receptors to the central nervous system and have cell bodies in the dorsal root or cranial sensory ganglia.^[4]^ Both neurons also differ in their target interfaces, with motor neurons forming specialized neuromuscular junctions and sensory neurons terminating in diverse sensory endings. These differences impose distinct requirements on biomaterial properties such as surface chemistry, mechanical compliance, and guidance cues.^[5]^ Moreover, sensory neurons generally exhibit a higher intrinsic regenerative capacity, grow faster, and are more robust than motor neurons, an important consideration for the development of nerve conduits and pro-regenerative material systems intended to promote selective or functional recovery following injury.^[3,6,7]^

As both cell types behave differently and have distinct needs, individual suitable materials, coatings, and scaffold architectures need to be developed, to mimic the matrix requirements, involving physical, biological, and mechanical characteristics.^[8]^ Hydrogel-based biomaterials are a promising approach for neural tissue engineering, as their mechanical and biochemical properties can be tailored to mimic the neural microenvironment and promote neural adhesion and neurite outgrowth in a three-dimensional (3D) culture.^[9]^ The materials should demonstrate biocompatibility and controlled degradation to enable neuronal cell adhesion, growth, differentiation, and the production of a native-like extracellular matrix (ECM).^[10]^ In addition to the biomaterial characteristics, the cell source is an important factor in establishing relevant neuronal tissue models or test biomaterials for regenerative therapies. For developmental studies or dense neuronal tissue formation, the use of stem cells is highly beneficial due to their high proliferation rate and differentiation potential. On the one hand, since stem cells require a large number of cell-cell contacts to maintain pluripotency and proliferation, the biomaterials used need to enable these interactions, while also allowing for dynamic self-organization during specific differentiation. On the other hand, the formation of dense tissue constructs requires sufficient nutrient and oxygen transfer as well as waste product removal; this could be facilitated through the biomaterial matrix until a perfusable vasculature is formed. An approach that fulfills both requirements are microporous annealed particle (MAP) scaffolds, as they provide large and variable pore sizes ranging from tens to hundreds of micrometers. MAP scaffolds are bottom-up-produced 3D microgel assemblies where spherical^[11]^ or rod-shaped^[12,13]^ microgels can be annealed via chemical interlinking or cell-induced interlinking.^[14]^ MAP scaffolds exhibit porosity with multi-cell-scaled macropores compared to nanoporous conventional bulk hydrogels. This allows for better cell infiltration and migration, while facilitating enhanced cell-cell communication, while the microgels can function as a pseudo-vasculature enabling diffusion of oxygen, nutrients, and other bioactive factors.^[11]^

To ensure neuronal cell growth in MAP scaffolds, the optimal biofunctionalization strategy needs to be identified. A large variety of protein and peptide coatings can be utilized for biomaterial functionalization to promote and support neuronal differentiation, interaction, and outgrowth.^[15]^ On a protein level, laminin (LN) is a promising candidate to support motor and sensory neuron interactions, as it promotes neurite outgrowth and supports neuronal differentiation,^[16,17]^ fibronectin (FN), an adhesive ECM protein, supports cell attachment and neurons by enhancing neurite outgrowth,^[18]^ and collagen IV (CollIV) coatings enhance neurite attachment and stabilize neurite-bearing phenotypes by extending collagen-binding sites for neuronal integrins.^[19]^ On a peptide level, isoleucine-lysine-valine-alanine-valine (IKVAV), a LN fragment, is known to enable neurite attachment and outgrowth by mimicking an active motif of the full-length protein. A similar effect is demonstrated for RGD, which is part of a variety of native ECM proteins^[20]^ and represents an integrin binding motif. In the context of iPSCs, the protein vitronectin (VTN) is abundantly used, but is comparably less studied regarding neuronal growth. However, it has been shown to support sensory neuron attachment and is likely to also support other neuronal subtypes, as it also contains the widely investigated attachment motif RGD.^[21]^ Tenascin-C (TN-C) is an ECM glycoprotein expressed during neural development that interacts with neuronal receptors in motor and sensory pathways. It plays a key role in regulating cell migration within the developing central nervous system and can modulate neurite outgrowth and cell adhesion.^[22]^ Overall, these proteins and peptides have shown positive results across both cell types, yet a direct comparison within the same experimental framework has not been investigated

In this study, we show that sensory and motor neurospheres can both be reproducibly differentiated and matured using a macroporous microgel-based scaffold platform. Synthetic PEG microgels are used as they are inherently bioinert, which allows us to test the influence of different biological coatings. The interaction of sensory and motor neurons with microgels presenting different biofunctionalization molecules was investigated, including LN, FN, CollIV, IKVAV, RGD, VTN, and TN-C. While pre-made precursor motor neurospheres demonstrated interaction only with VTN- and LN-coated microgels, the sensory neuron precursor spheres displayed interactions with all tested conditions. After the incorporation of microgels and further differentiation and maturation, neuronal spheroids were formed. The incorporation of microgels was quantitatively analyzed by visualizing the increase in spheroid area and measuring the increase of incorporated microgels. Furthermore, we generated iPSC-microgel scaffolds by leveraging iPSC cell-material interactions, subsequently differentiating the formed cellular scaffolds into motor or sensory neuronal lineages. We observed the strongest iPSC-material interaction with VTN-coated microgels, while LN-coated microgels still showed a strong interaction with iPSCs, leading to the formation of 3D cellular scaffolds, but with reduced microgel incorporation. The IKVAV-functionalized microgels showed no interaction with iPSCs and began interacting with the material only after 5 days of sensory neuron differentiation, indicating that the neuronal cell lineage was required for the interaction. We observed significant differences in growth kinetics between spheroids formed and differentiated directly from iPSCs and those formed from iPSC-derived precursor spheres, as well as between sensory and motor neuronal lineages, revealing the necessity of examining these processes to increase the impact for translational research. The herein reported material platform enabled the mechanistic study to test the influence of material biofunctionalizations on different neuronal subtypes separately, thereby posing great potential to further elucidate material optimizations for nerve regeneration and repair, which could be particularly valuable for regenerative SCI research.

## 2. Results and Discussion

### 2.1. Developing a microgel-based sensory and motor neurosphere platform

To create different neurosphere models, precursor motor or sensory neuron spheres or single iPSCs were cultured together with biofunctionalized PEG-microgels for 26 days in their corresponding differentiation media (Figure 1). The first experimental setup began with iPSC-derived embryoid bodies, formed in u-bottom well plates and differentiated for 5 or 7 days towards either the sensory or the motor neuronal lineage, respectively. The resulting precursor neuron spheres were subsequently cultured with PEG-microgels with different biofunctionalization molecules including: VTN, LN, IKVAV, CollIV, FN, RGD, and TN-C to evaluate cell-material interactions and the resulting cell-guided scaffold formation. The second experimental setup used a system previously developed in our group that combines iPSCs with differently VTN-functionalized PEG-microgels to create a stable self-formed cellular scaffold within 48 hours while maintaining stem cell pluripotency.^[23]^ Here, we investigated whether this is also possible with other coatings, like LN and IKVAV, and verified iPSC-induced cellular scaffold formation during 26 days of culture. After 2 days the resulting iPSC-microgel scaffolds were subsequently differentiated towards either the motor or the sensory neuron lineage following the protocol applied in the first setup. In both experimental setups, commercially available differentiation media were used to demonstrate the ease and user-friendliness of this platform, being accessible to a wide range of researchers and laboratories.

**Figure 1:**
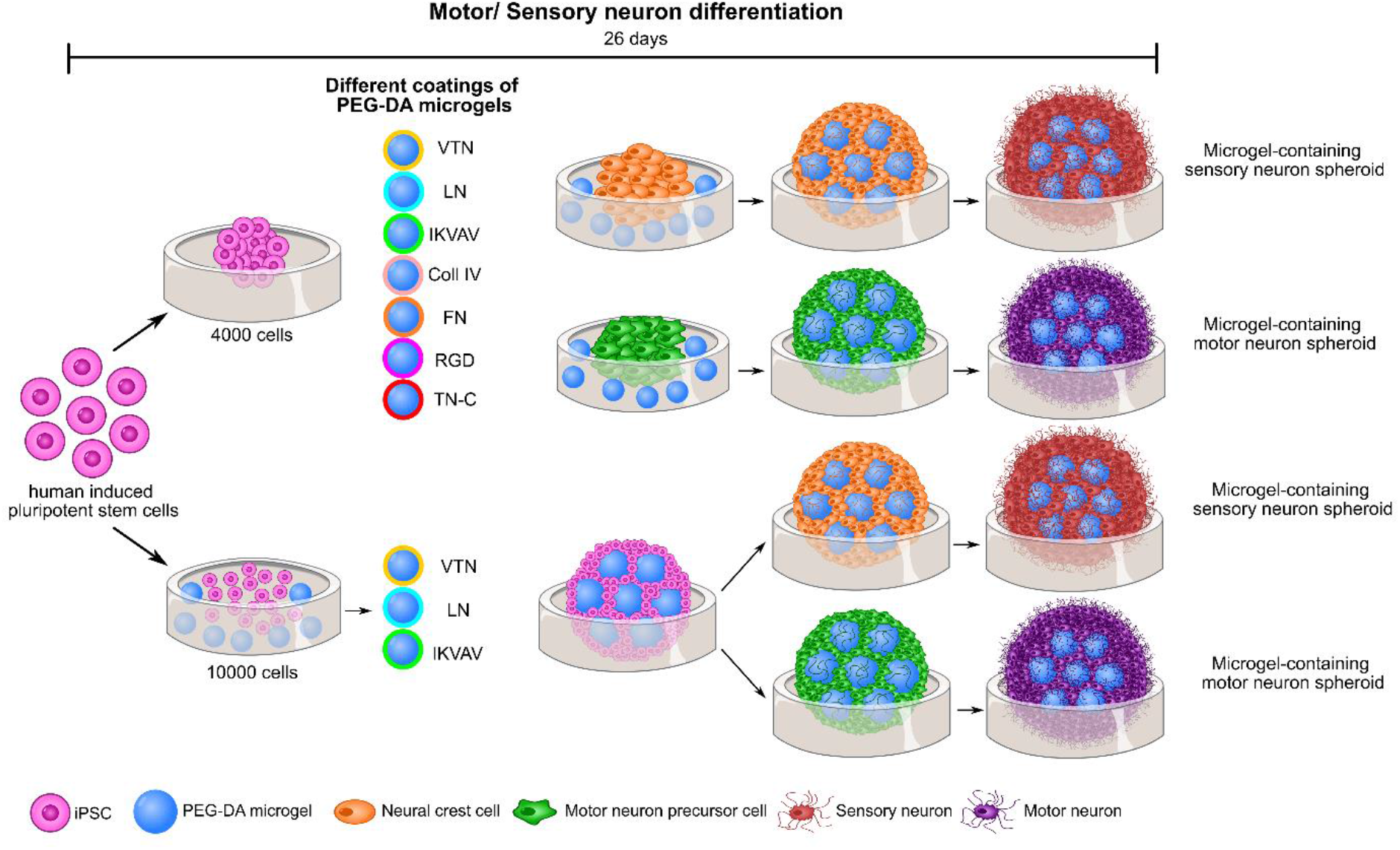
Schematic of the sensory and motor neurosphere differentiation. Precursor motor or sensory spheres or single iPSCs were cultured together with different biofunctionalized PEG-microgels for 26 days in their corresponding media.

In the initial approach, 5-day (sensory) or 7-day (motor) differentiated iPSC-derived precursor motor and sensory neuron spheres were combined with VTN-coated microgels. Different sizes of precursor spheres made with 2000, 4000, or 8000 cells were tested. It was evident that sensory neuron spheres began to incorporate the microgels during the first day of co-culture, while after three days, all the microgels were included in the spheroid (Figure SI1, bottom). This was observed for all different sensory spheroid sizes. Motor neuron precursor spheroids interacted less with the microgels (Figure SI1, above). After 1 day, no incorporation was visible, while after 3 days, slight incorporation of the microgels into the periphery of the spheroids was visible. This peripheral incorporation could have been caused by a slight interaction between the cells and the microgels. This indicates the need to further optimize the biomaterial platform to enable a strong motor neuron-material interaction. Furthermore, these results underlined the distinct differences in cell-material interactions between sensory and motor neuron precursor spheroids.

To further compare the differentiation and maturation of the different neuronal lineages over the course of 26 days, both precursor spheres were combined with the microgels at day 5 or 7 and further differentiated. Furthermore, the medium spheroid size, made with 4000 cells, was chosen for further experiments, in which different biofunctionalizations were tested to evaluate their compatibility with both neuron types.

### 2.2 Differentiation and maturation of precursor sensory neurospheres containing microgels

For sensory neurosphere formation, iPSCs were pre-differentiated for five days alone to ensure a neural crest path. This was followed by the incorporation of microgels with different biofunctionalizations (VTN, LN, IKVAV, CollIV, FN, RGD, and TN-C) at a ratio of 1 microgel per 100 cells to achieve microgel-containing spheroids; this was followed by further differentiation and maturation towards the sensory neuronal lineage (Figure 2A). Figure 2B shows the sensory neurosphere formation without microgels (only-cells, first row) and with LN-coated microgels (second row) over the time frame of 26 days. On the second day of culture, both spheroids appeared similar because no microgels were present. At day 6 the spheroids had incorporated the microgels, leading to an increase in spheroid size through microgel-induced volume increase and cell proliferation. The condition without microgels showed less size increase since, here, the increase was only caused by cell proliferation but not microgel incorporation. The same trend was observed for all other coatings, with a complete incorporation of the microgels within 24 to 48 hours, demonstrating strong cell-material interactions (Figure SI2). Another observation was the darkening of the spheroids throughout the 26 days culture time, indicating an increase in cellular scaffold density and thereby growth and compaction of the neuronal structures. At the endpoint (26 days), all sensory neurospheres expressed a uniform distribution of the general neuronal marker βIII-tubulin (TUJ1), as well as the specific sensory neuron differentiation marker peripherin (Figure 2C, SI3), demonstrating successful differentiation towards the sensory neuron lineage. The lack of other markers including the Schwann cell marker S100 in all conditions underlined the majority of the sensory neuron cell population (Figure SI4).

**Figure 2:**
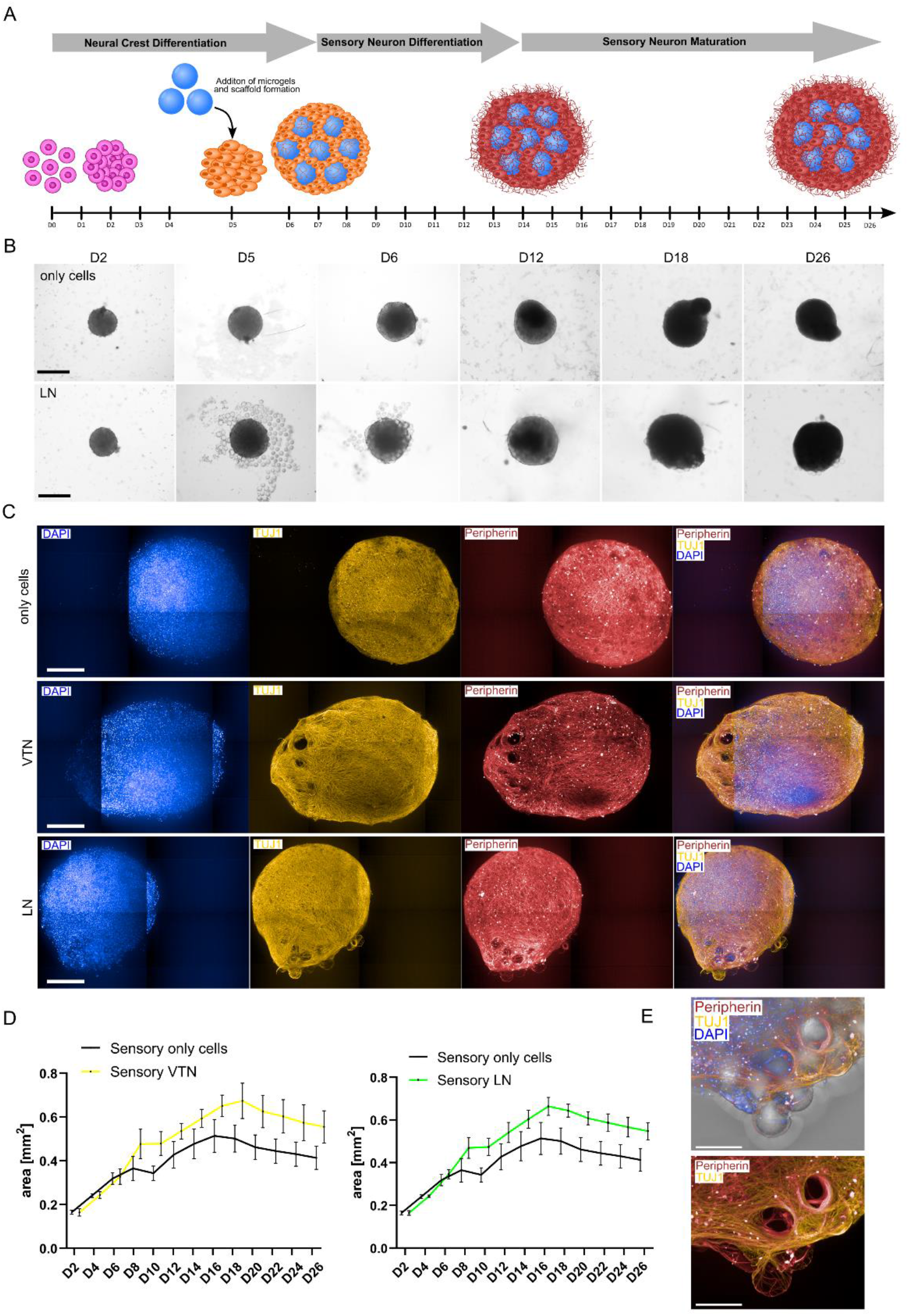
Sensory neurosphere differentiation and maturation from a precursor sensory neuron spheres. A. Schematic of the precursor sensory neuron sphere formation starting from single iPSC cells. After the addition of differently coated microgels, the spheroids differentiated and matured until day 26. B. Formation of only-cell (above) and with LN-coated microgel (below) sensory spheroids over 26 days in brightfield; scale bar is 500 µm. C. Sensory neurospheres at day 26 with only-cells (above), VTN-(middle), or LN-(below) coated microgels stained for DAPI (blue), TUJ1 (yellow), peripherin (red), and their corresponding merged image; scale bar is 200 µm. D. Area analysis of the sensory neurospheres over 26 culturing days, comparing spheroids cultured with VTN-coated microgels (yellow) and only-cells (black, first graph) or with LN-coated microgels (green) and only-cells (black, second graph). E. Close-ups of sensory spheroids cultured with VTN-coated microgels and showing their interaction in peripherin (red), TUJ1 (yellow), DAPI (blue), brightfield; scale bar is 100 µm.

For all the coated microgel conditions (VTN, LN, IKVAV, CollIV, FN, RGD, and TN-C), the incorporation of the microgels was clearly visible (Figure 2C, second and third row, Figure SI3). Close interactions between neuronal cell bodies/neurites and the microgels were observed as the latter followed their shape (Figure 2E). Due to the incorporation of microgels, an increase in cellular scaffold area was observed, validating the presence of microgels throughout the spheroids (Figure 2D, SI5). The area increase was similar across all biofunctionalized microgel conditions, resulting in a maximum area of around 0.6 mm² ± 0.1 mm², indicating that sensory neurons could interact with all tested proteins/peptides. In the first 17-19 days, the cellular scaffold area increased, with a higher increase for the microgel containing spheroids. The area comparison at days 8 and 18 (Figure SI9) underlined quantitatively that sensory neurons interact with microgels similarly across all tested conditions, whereas the only-cell condition showed a significantly lower area due to the absence of microgel-related spheroid volume increase. Interestingly, there is a spheroid area decrease after 17-19 days for both spheroids with and without microgel incorporation, potentially reflecting increased cellular rearrangement and compaction accompanying the formation of a maturing neuronal network,^[24,25]^. The observed decrease in spheroid area may also be associated with progressive neurite outgrowth and bundling, as growing neurites can generate traction forces that could contribute to the compaction.^[26]^ Additionally, for sensory neurons in late-stage differentiation, the ECM could remodel the structure, altering the content of highly hydrated ECM components, which could cause water retention and thereby spheroid size.^[25,27,28]^

These results showed the compatibility of the microgel platform with sensory neuron differentiation, maturation, and material interaction, proposing it as a suitable system for mechanistic studies.

### 2.3 Differentiation and maturation of motor neurospheres starting from precursor motor neuron spheres and microgels

Similar to the sensory neurospheres, the motor neurospheres also started from iPSC-derived precursor motor neuron spheres. The same number of microgels at a ratio of 1 microgel per 100 cells with the same coatings was used for the differentiation and maturation (Figure 3A). When the microgels were added, the precursor motor neurons demonstrated little to no material interaction, as visualized with brightfield imaging (Figure 3B, SI6). VTN-coated microgels showed minimal interactions at day 6, while LN, CollIV, and FN-coated microgels were slightly incorporated in the spheroid periphery at day 12. All other coatings promoted no interactions with the motor neurons. The delay in attachment could be explained by the stage-dependent changes in precursor cells, as early aggregates are dominated by cell-cell adhesions, whereas during maturation, integrin expression profiles, matrix adhesion, and neurite outgrowth can lead to interactions with particles on the spheres’ surface.^[29]^ The earlier interaction with VTN-coated microgels supported this hypothesis, as it is a protein known to interact strongly with iPSCs and neurons.^[30]^ Since during culture, non-incorporated microgels are reduced through media exchange, we here investigate the cell-material interaction of motor neurospheres in a precursor state.

**Figure 3:**
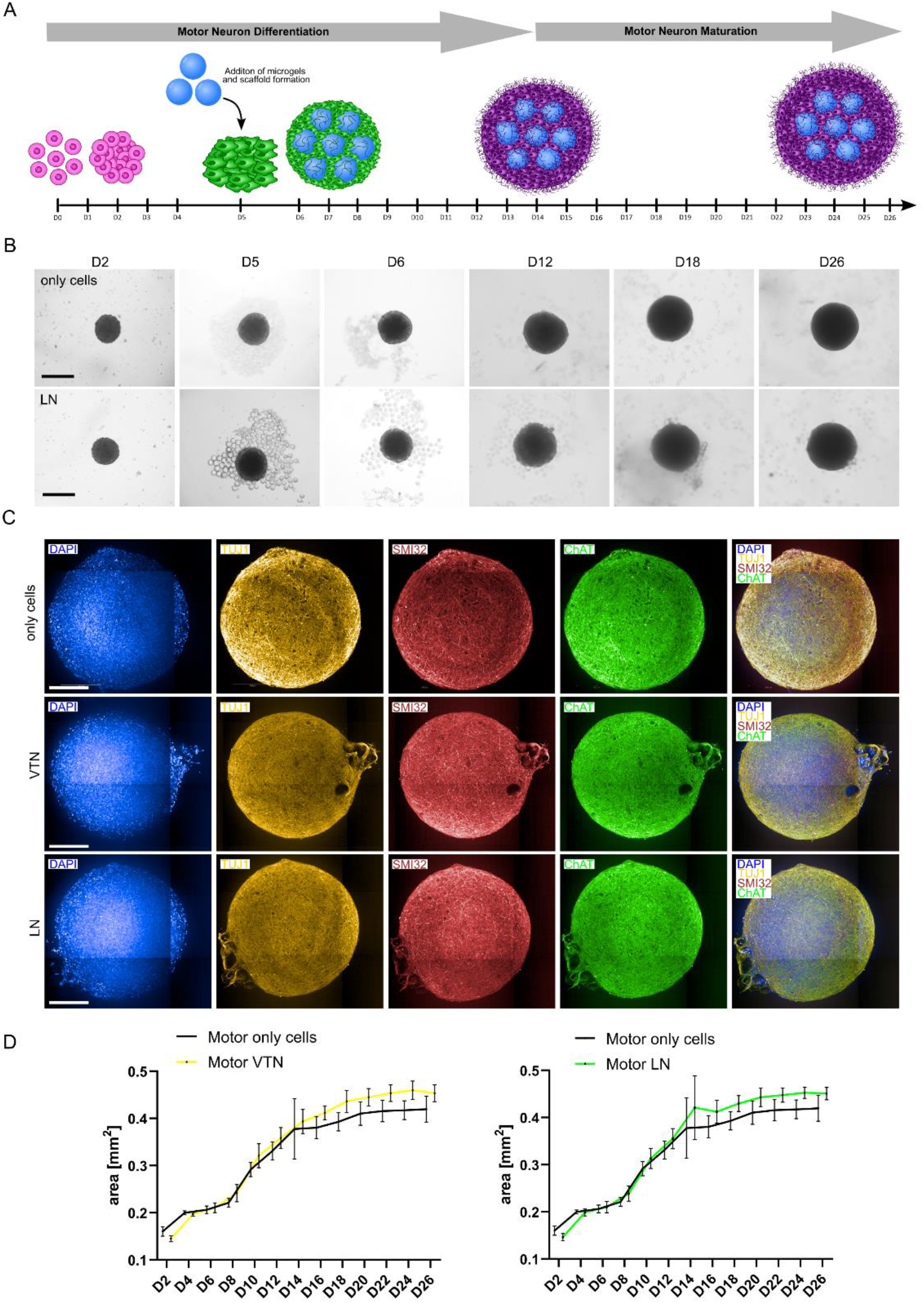
Motor neurosphere differentiation and maturation from a precursor motor neuron sphere. A. Schematic of the precursor motor neuron sphere formation starting from single iPSC cells. After the addition of differently coated microgels, the spheroids differentiated and matured until day 26; B. Formation of only-cell (above) and with LN-coated microgel (below) motor spheroids over 26 days in brightfield; scale bar is 500 µm. C. Motor neurospheres at day 26 with only-cells (above), VTN-(middle), or LN-(below) coated microgels stained for DAPI (blue), TUJ1 (yellow), SMI32 (red), ChAT (green), and their corresponding merged image; scale bar is 200 µm. D. Area analysis of the motor neurospheres over 26 culturing days, comparing spheroids cultured with VTN-coated microgels (yellow) and only-cells (black, first graph) or with LN-coated microgels (green) and only-cells (black, second graph).

Similar to the sensory neurons, the spheroids became darker and denser over time. After 26 days, all spheroids showed a uniform distribution of the neuronal marker TUJ1, as well as the specific motor neuron marker SMI32 that binds non-phosphorylated neurofilament H, which defines mature projection neurons, like motor neurons in the spinal cord,^[31]^ and choline acetyltransferase (ChAT), which functions as a marker for cholinergic neurons^[32]^ (Figure 3C, SI7). Similar to the brightfield, it was visible that the spheroids were attached to a small number of microgels. The area analysis showed a slightly larger area size for the microgel-containing conditions compared to the only-cell spheroids (Figure 3D, SI8). We hypothesized that the increase in area correlates with cellular growth kinetics, influenced by the different protein or peptide coatings presented, even without strong cell-material interactions and microgel incorporation. All conditions showed the same trend: a slow size increase in the first ∼8 days of culture, followed by an increased linear growth until ∼day 16, ending in a slow balanced growth until 26 days. The area growth curves verified that the motor neuron precursor spheres did not incorporate the microgels, as their area showed no significant difference from the only-cell conditions at days 8 and 18 (Figure SI9, left row). Compared to the sensory neurospheres, which compact due to maturation-related neurite bundling, the motor neurospheres did not decrease in size over the 26 culturing days. This could indicate a very dense initial cell packing in the spheres with strong cell-cell interactions, leading to no further compaction. Additionally, motor neurons show less neurite outgrowth than sensory neurons, leading to less mechanical forces exerted by the lower number of neurites wrapping around the microgels. To conclude, we observed only a slight positive effect of full-length proteins on motor neuron growth, despite weak adhesion, indicating the specificity of the system to changes in signaling. Furthermore, we underlined the importance of comparing sensory and motor neuron systems mechanistically in biomaterial design, due to their intrinsic material interaction capabilities.

### 2.4 Sensory neurosphere formation from iPSC-microgel cellular scaffolds

As the motor neuron precursor spheroids appeared to have difficulties incorporating the microgels, we decided to start directly from single iPSCs to enable microgel-containing motor neurospheres. Firstly, we tested the formation of sensory neurospheres, followed by motor neurospheres (see 2.5). Therefore, 10,000 iPSCs were cultured together with around 80 microgels, in the same cell-to-microgel ratio as in the neuron precursor experiments, for 2 days in iPSC maintenance media to allow the formation of pluripotent iPSC-microgel cellular scaffolds, followed by differentiation into sensory neurospheres for 26 days (Figure 4A). Three different coatings were tested: VTN, LN, and IKVAV. Based on the known interactions between iPSCs and VTN and LN, we compared the known iPSC niche ECM protein VTN with the neurite growth- and differentiation-promoting ECM protein LN. IKVAV was included as a functionalization to test whether iPSCs can interact with this short neurite growth-promoting peptide sequence from LN, which is known for neuron adhesion, but has not been reported regarding iPSC interaction.

**Figure 4:**
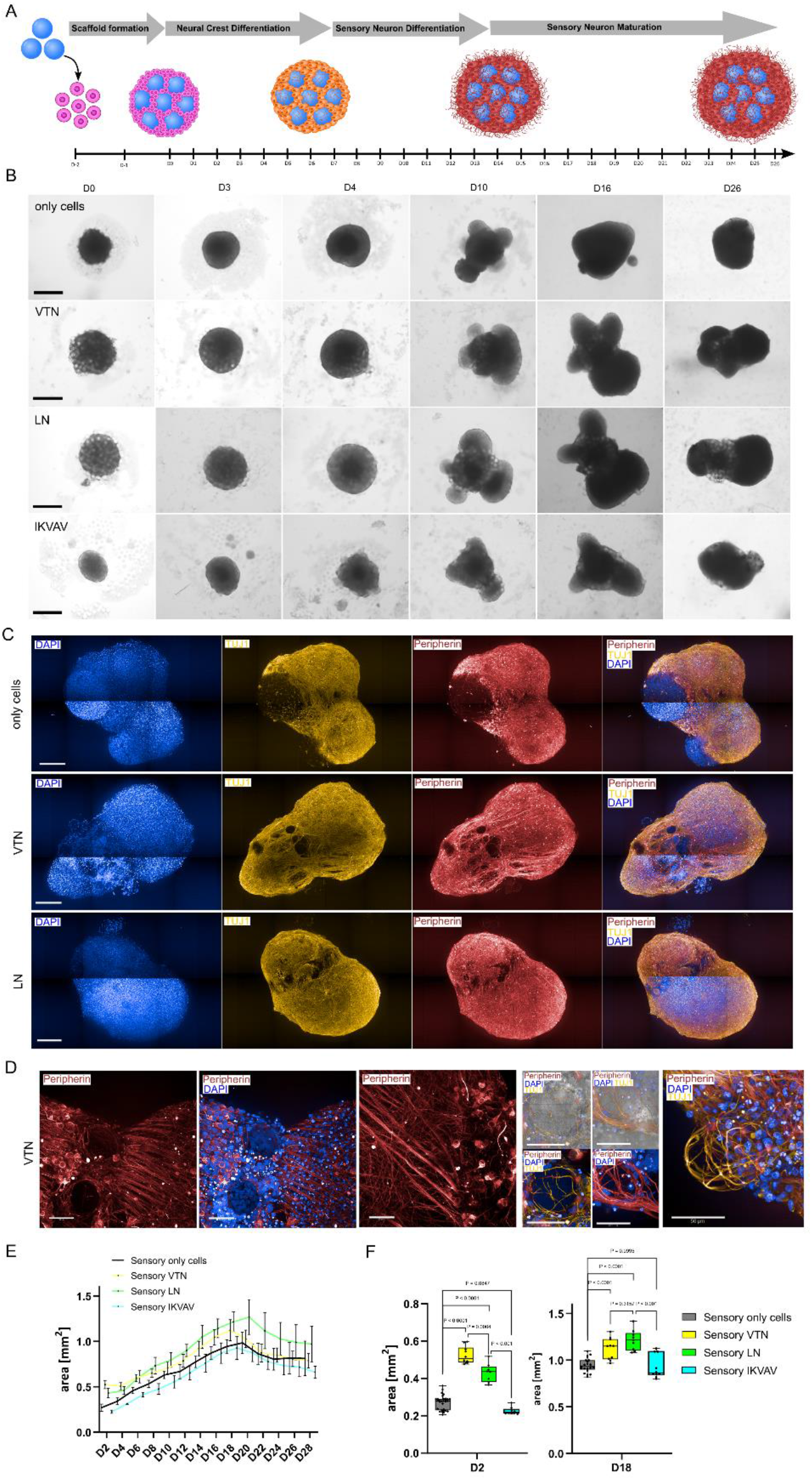
Sensory neurosphere differentiation and maturation from single iPSCs. A. Schematic of the sensory neurosphere formation starting from single iPSCs together with differently coated microgels. The spheroids differentiated and matured until day 26; B. Formation of only-cell (first row), with VTN-(second row), LN-(third row), and with IKVAV-(fourth row) coated microgel sensory spheroids over 26 days in brightfield; scale bar is 500 µm. C. Sensory neurospheres at day 26 with only-cells (above), VTN-(middle), or LN-(below) coated microgels stained for DAPI (blue), TUJ1 (yellow), peripherin (red), and their corresponding merged image; scale bar is 200 µm. D. Close-ups of sensory spheroids cultured with VTN-coated microgels and showing their interaction in peripherin (red), TUJ1 (yellow), DAPI (blue), brightfield; scale bar is 100 µm (first 3 images) and 50 µm (last 5 images). E. Area analysis of the sensory neurospheres over 26 culturing days, comparing spheroids cultured with VTN-coated microgels (yellow), with LN-coated microgels (green), IKVAV-coated microgels (cyan), and only-cells (black). F. Area comparison of the sensory neurospheres after 2 and 18 culturing days, comparing spheroids cultured with VTN-coated microgels (yellow), with LN-coated microgels (green), IKVAV-coated microgels (cyan), and only-cells (black).

Here, we observed higher growth in the spheroids compared to the precursor neuron condition due to the higher initial cell number (10,000 vs 4,000 cells) and the high proliferation rate of iPSCs. Interestingly, VTN- and LN-coated microgels were all incorporated after three days of culture (Figure 4B, second and third row). For the IKVAV-coated microgels, iPSCs did not demonstrate cell-material interactions and microgel incorporation, which is likely due to the weak interaction between IKVAV and iPSCs (Figure 4B, fourth row). After 5 days of differentiation (thus day 7 of culture) towards the sensory neuronal lineage, the neuronal crest spheres started to incorporate IKVAV-coated microgels, demonstrating the interaction between neuronal precursor cells and IKVAV. Sensory neuron differentiation can cause a shift in integrin expression profiles, introducing the IKVAV-binding capacity, since interactions with IKVAV are known to be highly dependent on sufficient β1-integrin expression and clustering.^[33–35]^ These results underlined the importance of cell-type-specific biofunctionalization and the time point of material addition. Around day 10, all spheroids, regardless of the microgel coating and similar to the condition without microgels, started to form multiple uneven outgrowth zones, which increased in size up to day 16, before forming a dense, rounder structure until day 26. Comparable to the precursor experiments, we observed a uniform expression of TUJ1 and peripherin (Figure 4C, SI10A) across all tested microgel-containing conditions. In contrast, the only-cell spheroid showed a less uniform signal of neuronal markers, with areas without differentiation towards the neuronal lineage (Figure 4C, first row). Furthermore, we observed in the sensory neurospheres with VTN-coated microgels, the formation of thick peripherin-positive fiber bundles (Figure 4D), indicating an increased maturation and improved differentiation beginning from iPSC-microgel scaffolds compared to precursor sensory neurospheres. Additionally, the spheroids did not show expression of S100, indicating that the neuronal marker-negative area in the only-cell condition did not differentiate towards Schwann cells (Figure SI10B). The brightfield images at day 26 showed dark spots in certain spheroid regions (Figure SI11), especially in the only-cell and IKVAV-coated microgel conditions, which could indicate differentiation into melanocytes, as the black deposits observed are hypothesized to be the dark pigment melanin, which is stored inside the melanosomes of melanocytes.^[36,37]^ This hypothesis is further strengthened by the fact that melanocyte development takes place from neural crest precursors, thereby sharing the same precursor state as sensory neurons.^[38]^ The VTN- or LN-coated microgel conditions did not demonstrate the formation of these dark areas. The area growth was quantified for VTN- and LN-coated microgels compared to the only-cell condition, demonstrating microgel incorporation (Figure 4E). The IKVAV-coated microgels were not incorporated, which resulted in an initially smaller area, even compared to the only-cell condition, indicating a negative influence of IKVAV on the iPSC proliferation. Over time, spheroid growth was observed across all conditions. Interestingly, while the VTN-microgel condition initially had the largest area (2 days), the LN condition showed an increased growth, resulting in the largest spheroid area from day 6 until the end of culture. The measured area of the VTN condition on day 2 was 0.52 ± 0.05 mm^2^, compared to the LN condition with 0.43 ± 0.05 mm^2^. Comparing the area sizes at a later stage, the LN condition grew to an area of 1.22 ± 0.11 mm^2^ on day 18, equaling a size increase with around 184% since day 2 compared to 115% for the VTN condition, resulting in an area of 1.12 ± 0.12 mm^2^. The IKVAV condition remained with the smallest spheroid area throughout the first 18 days of culture, even though the microgels were incorporated partly after 6 days of culture (Figure 4F). After 18 days, the size of the IKVAV condition increased to a comparable size with the only-cell condition. The IKVAV-coated microgel-containing spheroids grew from 0.23 ± 0.02 mm^2^ (day 2) to 0.93 ± 0.13 mm^2^ (day 18), demonstrating an area increase, with 304% compared to the only-cell control, with 0.27 ± 0.04 mm^2^ (day 2) to 0.94 ± 0.08 mm^2^ (day 18), equaling an increase of 248%. Concluding, we observe a higher increase in area over 16 days for the IKVAV condition and the only-cell condition, compared to the VTN and LN conditions. This could be explained by the positive ECM-mediated effects of integrin binding to VTN and LN, resulting in faster differentiation and maturation of the sensory neuron lineage, thereby decreasing the time in the highly proliferative precursor stage, while increasing the time in the low-proliferative and stagnating maturation stage. The slightly higher area increase rate for the IKVAV-coated microgel condition compared to the only-cell-control could be explained through the additional incorporation of the microgels during neuron differentiation observed earlier, which resulted in a volume and, thereby, area increase. These assumptions are further supported through the thick fiber formation in sensory neurospheres with VTN-coated microgels, elucidating a further maturation state compared to the other conditions. These fiber bundles were oriented in a single direction throughout the construct and exhibited a thickness of up to 50 µm, indicating mature structural orientation and sensory neuron maturation.

### 2.5 Motor neurosphere formation from iPSC-microgel cellular scaffolds

We further tested the use of the iPSC-microgel cellular scaffold platform for the formation of motor neurospheres, enabling the incorporation of microgels into the spheroids in contrast to the precursor motor neuron setup described in 2.3. To compare the results with the previously tested sensory neuron differentiation, 10,000 iPSCs were combined with differently biofunctionalized microgels (VTN, LN, IKVAV), and the resulting spheroids were differentiated and matured for 26 days in culture (Figure 5A). The incorporation of microgels directly by the iPSCs and thereby the cellular scaffold formation before the induction of motor neuronal differentiation showed the same trend as the sensory neurospheres formed by an iPSC- microgel cellular scaffold. While the iPSCs exhibited strong cell-material interactions with VTN- and LN-coated microgels, they did not interact with IKVAV-coated microgels (Figure 5B). Unlike sensory neuron precursors, motor neuron precursors did not begin incorporating the IKVAV-coated microgels at day 5, highlighting the differences between these neuronal subtypes. After 26 days, we observed the expression of the neuronal marker TUJ1 and the specific motor neuron markers SMI32 and ChAT across all conditions (Figure 5C, SI12). In line with the observations made for sensory neurospheres, we observed a uniform distribution of motor neuronal marker expression across all microgel-containing conditions, whereas the only-cell-only condition showed small outgrowth regions without neuronal markers. These outgrowth regions were spanned by neurites but contained no neuronal cell bodies (Figure 5C). Furthermore, we observed the formation of thick SMI32-positive fibers in conditions with the VTN- and LN-coated microgels, indicating a further maturation state of the motor neurons by an increase in neurite complexity.^[39]^ The trend in motor neurosphere size was similar to the observed trend for the sensory neurospheres (Figure 5D). Initially, the VTN-coated condition demonstrated the largest spheroid area due to the preferred interaction between iPSCs and the VTN. During culture time, however, the LN condition demonstrated the largest spheroid area, indicating the beneficial effect of the LN interaction on motor neuron growth. The IKVAV- coated microgels, on the other hand, resulted in smaller spheroids, comparable to the only-cell condition. This could be explained since the iPSCs did not demonstrate interactions with the IKVAV-coated microgels, as observed before, but contrary to the sensory neuron precursors, the motor neuron precursors did not start incorporation of the microgels during differentiation. All conditions reached a plateau after 26 days. These trends were further visualized and statistically analyzed in the area comparison on days 2 and 18 (Figure 5E), where first VTN was significantly higher than LN on day 2, and slightly lower on day 18. The only-cell and IKVAV conditions, while being non-significantly different from each other, were significantly smaller than the VTN and LN conditions. The cellular scaffolds formed at day 2 with VTN-coated microgels demonstrated the largest area with 0.51 ± 0.8 mm^2^, followed by the LN condition with 0.38 ± 0.07 mm^2^. With no microgel incorporation observed, the IKVAV condition demonstrated a significantly smaller area with 0.23 ± 0.01 mm^2^, comparable to the only-cell control with an area of 0.26 ± 0.03 mm^2^. Further, we evaluated the spheroid size on day 18 of culture, observing an overtake of the LN-coated microgel condition at later time points, presenting the largest size with 0.66 ± 0.06 mm^2^, compared to the VTN condition with an area of 0.60 ± 0.05 mm^2^, indicating the positive influence of LN on neurite extension and maturation, observed as well in the sensory neuron lineage. The area of the IKVAV-coated microgel condition increased to 0.52 ± 0.06 mm^2^, and the only-cell-condition had an area of 0.53 ± 0.09 mm^2^. Between 2 and 18 days, we saw an increase in the area for the VTN condition with 17%, for the LN condition with 74%, for the IKVAV condition with 126%, and for the only-cell condition with 104%. The trend is thereby identical to the one observed for the sensory neuron lineage, with a striking difference in the general percentages of area increase, for example, around a 2.5-fold higher percentage in the LN condition.

**Figure 5:**
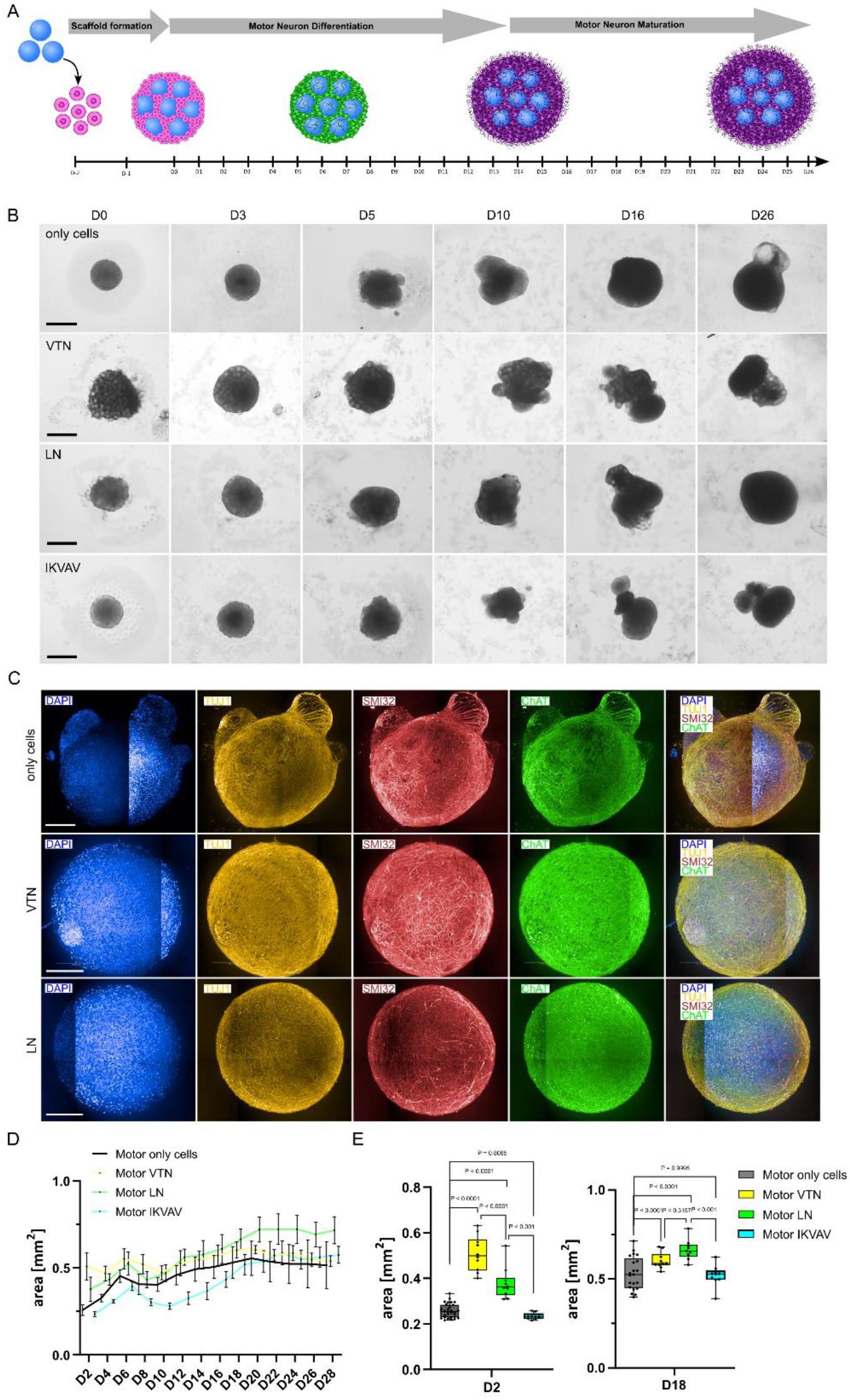
Motor neurosphere differentiation and maturation from single iPSCs. A. Schematic of the motor neurosphere formation starting from single iPSCs together with differently coated microgels. The spheroids differentiated and matured until day 26; B. Formation of only-cell (first row), with VTN- (second row), LN- (third row), and with IKVAV-coated microgel (fourth row) motor spheroids over 26 days in brightfield; scale bar is 500 µm. C. Immunostained images of motor neurospheres at day 26 with only-cells (above), VTN-(middle), or LN-(below) coated microgels stained for DAPI (blue), TUJ1 (yellow), SMI32 (red), ChAT (green), and their corresponding merged image; scale bar is 200 µm. D. Area analysis of the motor neurospheres over 26 culturing days, comparing spheroids cultured with VTN-coated microgels (yellow), with LN-coated microgels (green), IKVAV-coated microgels (cyan), and only-cells (black). E. Area comparison of the sensory neurospheres after 2 and 18 culturing days, comparing spheroids cultured with VTN-coated microgels (yellow), with LN-coated microgels (green), IKVAV-coated microgels (cyan), and only-cells (black).

The more homogeneous marker distribution in the microgel-containing spheroids supports the hypothesis of a positive influence of enhanced diffusion of differentiation factors from the media and the presence of bioactive molecules through microgel incorporation.

### 2.6 Assembloid formation of motor neuron and skeletal muscle spheroids

To investigate the functionality and maturity of our motor neurospheres, we cultured them together with a human skeletal muscle cell spheroid. For this, we first established the muscle spheroid. We combined RGD-coated rod-shaped microgels with single human skeletal muscle cells to form skeletal muscle/microgel spheroids. Rod-shaped instead of spherical microgels were chosen for the muscle spheroids due to the reported beneficial effect of anisotropy on muscle cell morphology and myogenic marker expression.^[40]^ After one day, the muscle cells adhered to the rods and were cultured in differentiation media until day 7. After this, they were added to the 26-day-old motor neurosphere and cultured together for an additional 7-day period (Figure 6A). We observed the infiltration of thick TUJ1-positive neurites into the skeletal muscle spheroid, demonstrating the interaction between the motor neurons and the skeletal muscle cells (Figure 6B). While the neurites inside the motor neurosphere were positive for SMI32, the extended neurites towards the muscle spheroid did not express SMI32. This indicated the ability of the motor neurons to transition between a more mature motor neuron state and an infiltrating, more “target searching” state, in case of contact with muscle cells, potentially demonstrating motor neuron functionality. (Figure 6C, D, E). We further compared the infiltration of motor neurons into muscle spheroids with the infiltration into sensory neurospheres. We did not observe the thick neurite bundles observed for muscle spheroid infiltration, further underlining the ability of the motor neurons to sense and interact differently with muscle cells (Figure SI13). These different assembloid formations showed that our motor neurospheres are mature enough to show initial interactions with the muscle spheroids, demonstrating a promising approach for future use of these spheroids in neuromuscular junction research.

**Figure 6:**
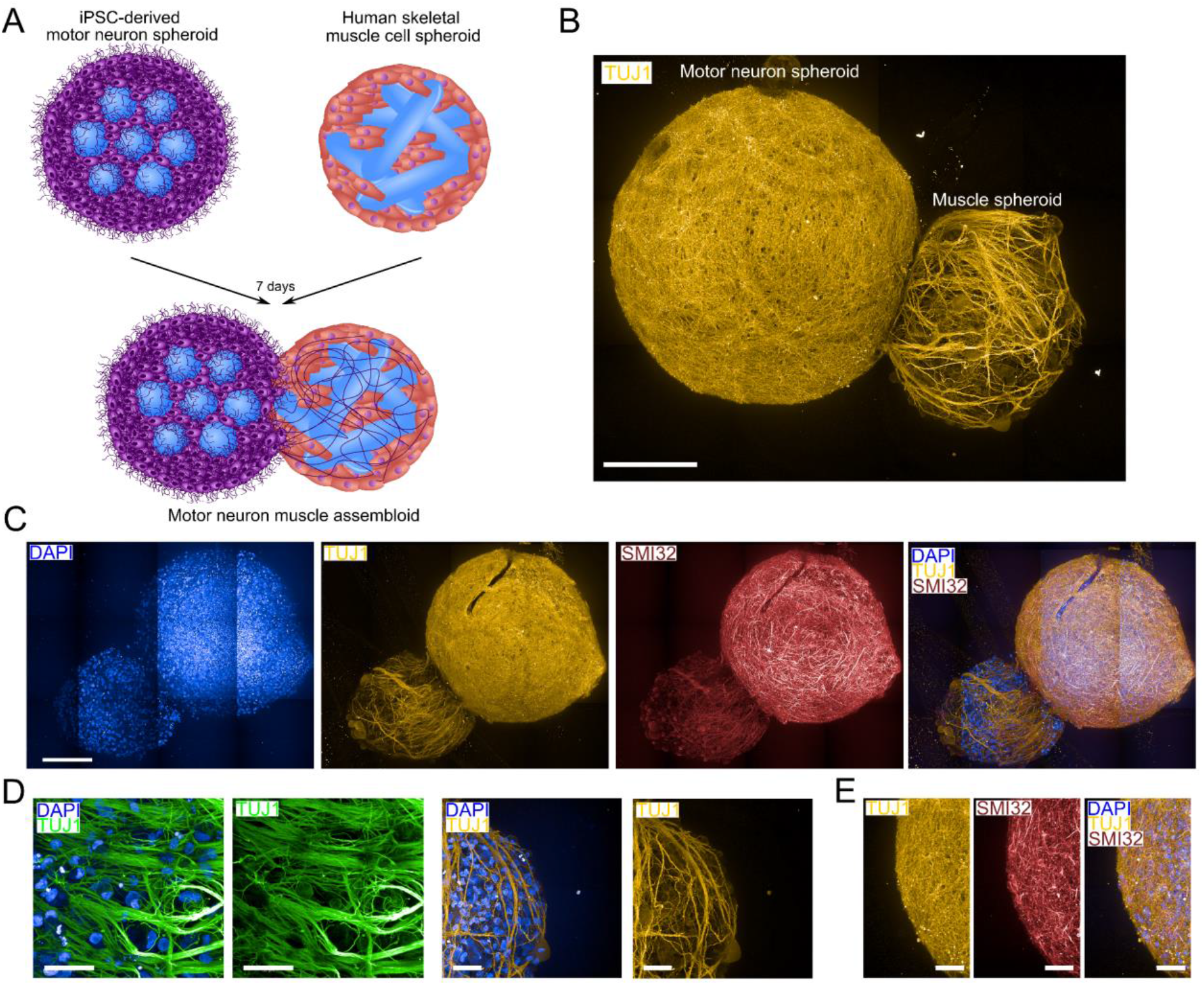
Assembloid of motor neurospheres with skeletal muscle spheroids. A. Schematic of the motor neurosphere and skeletal muscle assembloid formation. B, C. Immunostained images of motor neuron muscle assembloids at day 32 stained for DAPI (blue), TUJ1 (yellow), SMI32 (red) and their corresponding merged image; scale bar is 200 µm; D. Close-up of the muscle spheroid with motor neurons and muscle cells stained for DAPI (blue) and neurites stained with TUJ1 (green (left), yellow (right)); scale bar is 50 µm; E. Close-up of the motor neurosphere stained for TUJ1 (yellow), SMI32 (red) and their corresponding merged image; scale bar is 50 µm.

## 3. Conclusion and Outlook

We established two different biomaterial-based systems, starting either from precursor neurospheres (sensory or motor) or naïve iPSCs, which were combined with differently coated PEG-based microgels to form 3D cellular scaffolds. The systems were further differentiated and matured into sensory or motor neural lineages. By comparing seven different ECM-associated microgel coatings (VTN, LN, CollIV, FN, TN-C, RGD, and IKVAV) in the precursor systems and three different coatings (VTN, LN, and IKVAV) in the naïve iPSC cellular scaffold systems, we showed that cell-material interaction depends on neuronal subtype and differentiation stage, and that it is influenced by the time at which material is incorporated during differentiation. In particular, precursor sensory and motor neurospheres showed cell-material interactions to be lineage-dependent. Sensory neurospheres demonstrated strong cell-material interactions across the different microgel coatings and showed increases sizes (as indicated by larger areas) neurosphere. Motor neurons, on the other hand, incorporated fewer or no microgels, underlining their weaker cell-material interactivity. As expected, at a naïve stage, iPSCs strongly interacted with VTN- and LN-coated microgels, while IKVAV-coated microgels excreted no response. In contrast to the motor neuronal differentiation condition, the sensory neuron condition, showed incorporation of IKVAV-coated microgels at day 5, indicating that the sensory neuronal precursors were ready for interaction with the peptide already at this early stage. The motor neuron condition, however, did not demonstrate any interaction with the IKVAV-coated microgels, mirroring the observations made with the precursor motor neurospheres. The motor neurosphere cellular scaffold showed thick SMI32- positive neurites, while the sensory neurosphere ones formed peripherin-positive multi-neurite fiber bundles, which were not visible in the precursor neurospheres or the conditions without microgel incorporation. The potential functional relevance of motor neurospheres was indicated in neuromuscular assembloids with human skeletal muscle spheroids. The muscle spheroid was infiltrated by neurosphere neurites from the motor cellular scaffold in defined strong TUJ1- positive neurite bundles. In comparison, when paired with sensory neurspheres thick neurite bundles were missing, supporting target-specific response.

These findings emphasize that biomaterials exert lineage- and stage-specific responses that cannot be generalized across lineage and developmental contexts and present an initial step in understanding lineage-specific requirements in biomaterial design. For future experiments, electrophysical analysis, such as patch clamp or multielectrode (MEA) measurements, together with calcium imaging, could be performed to confirm neuronal network activity. This microgel-based platform proposes a useful method for systematically investigating the distinct biomaterial requirements of iPSC-derived motor and sensory neurons. Furthermore, the neurospheres have the potential to be combined into assembloids, enabling the study of neuromuscular interactions, sensory-motor circuits, or other tissue interfaces. Overall, this platform demonstrates a straightforward and easily adaptable methodology to test proteins, peptides, or peptide mimetics for their interaction with both motor and sensory neurons, providing a valuable tool for biomaterial development in SCI regeneration.

## 4. Experimental Section/Methods

### Cell culture

The iPSC line (UKAi009-A) was used in this study, and further information can be found in the Human Pluripotent Stem Cell Registry (hPSCreg).^[41]^ Maintenance cell culture of iPSCs was performed in tissue culture-treated 6-well plates coated with Vitronectin (Vitronectin XF^TM^ STEMCELL technologies^TM^, # 07180; 0.5 µg/cm^2^) in StemMACS™ iPS-Brew XF (Miltenyi biotec, 130-104-368). Cells used during this work were between passages 12 and 30.

Human skeletal muscle cells (passage 3-5, Promocell) were cultured in tissue culture flasks in Skeletal Muscle Cell Growth Medium (Promocell) with 1% AMB. For differentiation, Skeletal Muscle Cell Differentiation Medium (Promocell) was used for 6 days.

### Microgel preparation

Microgels with a diameter ∼ 80 µm were prepared after Klasen et al.^[23]^ Briefly, epoxy-functionalized microgels were prepared from aqueous solutions composed of polyethylene glycol diacrylate (PEG-DA, 15% w/w, Mn 700, Sigma-Aldrich), Lithium-Phenyl-2,4,6- trimethylbenzoylphosphinate (LAP, 1% w/w, Sigma-Aldrich) photoinitiator, and freshly distilled glycidyl methacrylate (GMA, 1% w/w, Sigma-Aldrich). As a control, unfunctionalized PEG microgels were produced analogously without the addition of GMA. The solution was stored in brown glass vials until use to protect from undesired light exposure. The continuous oil phase was composed of Novec™ HFE 7500 with Krytox™ 157 FSH surfactant (4.5% v/v, DuPont). Both solutions were transferred into a syringe (Hamilton Gastight® Series 1000, 10mL), and the syringe containing the dispersed phase was wrapped in aluminum foil. A 25G cannula was inserted into PTFE tubing (OD 0.9 mm, ID 0.4 mm, TECHLAB GmbH) and attached to the syringes. Residual air was removed, and the syringes were fixed in a syringe pump (Harvard Apparatus Pump 11 Elite). The other ends of the tubes, as well as an outlet tube (Polyethylene, OD 0.038 in ID 0.023 in, Instech Laboratories), were inserted into the dedicated inlet holes on the microfluidic device. The continuous phase was started at a flow rate of 1000 µL/h to prefill the collection reservoir. Afterwards, the dispersed phase was started at 1000 µL/h until droplet formation was visible. The flow rates were then adjusted to 3000 µL/h and 2000 µL/h for the continuous and dispersed phase, respectively. Droplet crosslinking was achieved by guiding the outlet tube through a self- constructed UV-LED set-up, as previously reported.^[12]^ Briefly, the UV set-up contains a fixed frame featuring an array of five LEDs (Starboard, Luminus SST-10-UV-A130, 365 nm) spaced at a distance of 23 mm, powered by a 24 V power supply, and operating at an irradiation dose of 30.36 mW/cm2. The microgels are crosslinked in-flow by fixing the outlet tube in a PDMS gasket connected to the frame. Purification of the microgels was achieved by multiple subsequent washing steps, as previously reported.^[42]^ In short, the excess continuous phase was pipetted off, and the microgels were sequentially washed with Novec™ HFE 7500, n-hexane + span 80 (1% v/v), n-hexane, IPA, and Milli-Q water. Each solvent was applied five times before proceeding to the next solvent. After the purification, the microgels were stored in Milli-Q water at 4 °C until use.

### Biopeptide- functionalization of microgels

Before functionalization with different biopeptides, the microgels, suspended in 1x PBS pH 7.4, were UV-sterilized for 2 hours and subsequently handled only under sterile conditions. Microgels were functionalized with the Vitronectin (VTN) XF™ solution (STEMCELL technologies^TM^), Laminin (LN) (Sigma Aldrich), Fibronectin (FN) (Sigma Aldrich), IKVAV (Ac-FKGGAAGGEEEEGIKVAV, Genscript), RGD (GRGDSPC, GPC Scientific), Collagen type IV (CollIV) (from human placenta, Merck), and Tenascin C (TN-C) (Acro Biosystems). Therefore, microgels were pelleted and resuspended in an equal volume of 1x PBS corresponding to the pellet volume (e.g., 100 µL microgel pellet + 100 µL 1x PBS). To this, 240 µL volumes of the VTN-solution at 250 µg/mL were added, resulting in a final amount of 60 µg VTN added. The other full-length proteins, LN, TN-C, FN, and CollIV, were added with 60 µg per 100 µL microgel pellet, respectively. The peptide motifs were added with 500 µg IKVAV or RGD for 100 µL of microgel pellet (RGD-stock 25 mg/mL; IKVAV-stock 5 mg/mL). The final solution was incubated overnight at 4 °C and for 20 minutes at 37 °C directly before use for experiments. The microgels stored in the coating solution were centrifuged (1 min, RT, 8000 rpm) and the coating solution was replaced with iPS-Brew media, in a ratio of 4:1 media to microgel pellet volume (e.g., 400 µL media + 100 µL microgel pellet). For the experiment, microgels were pipetted straight from the pellet. The respective microgel number added to the experiments using this microgel solution was calculated using a calibration curve.^[23]^

### iPSC-based precursor motor/sensory neuron sphere generation

Human induced pluripotent stem cells were cultured on VTN (STEMCELL technologies)- coated tissue culture plates. After reaching 70-80% confluency, 4.000 iPSCs were seeded in a 96- well ultra-low attachment U-bottom plate (Thermofisher scientific^TM^, 174925), filling up the media volume to 150 µL, and centrifuged for 4 min at 300 rcf using an Eppendorf Centrifuge 5810 R. Directly from day 0, we started using the motor and sensory neuron differentiation kits from Stemcell Technologies. In case of sensory neuron differentiation, we used the STEMdiff™ Neural Crest Differentiation Kit (Stemcell technologies, #08610) until day 7 of differentiation, followed by the STEMdiff™ Sensory Neuron Differentiation Kit (Stemcell technologies, #100-0341) until day 14 of differentiation. Afterwards, we matured the sensory neurons for an additional 12 days using the STEMdiff™ Sensory Neuron Maturation Kit (Stemcell technologies, #100-0684). The media was changed daily during the complete culture. For the motor neuron differentiation, we applied the STEMdiff™ Motor Neuron Differentiation Kit (Stemcell technologies, #100-0871) for the first two weeks, followed by the maturation phase of 12 days using the STEMdiff™ Motor Neuron Maturation Kit (Stemcell technologies, #100-0872). We also performed daily media changes. For both sensory and motor neuron protocols, we added 10 µM ROCK inhibitor (Y-27632 (Dihydrochloride), STEMCELL technologiesTM, # 72304) to the media composition for the first 24 hours of the experiment. The media composition and timepoints of media change were performed according to the protocols provided by the media manufacturer. The complete culture was performed in the Nunclon™ Sphera™ U-well 96-well plates (Thermofisher scientific^TM^, 174925). At day 5 of neural lineage differentiation, 0.25 µL of prepared microgel solution (prepared in the same media) was added to each precursor neuron sphere formed and the differentiation protocols were continued for 26 days.

### iPSC-microgel scaffold generation and motor/sensory neuronal differentiation

After reaching around 80% confluency, iPSCs were detached into single cells using ACCUTASE^TM^ (STEMCELL technologiesTM, # 07920) cell detachment solution. 0.625 µL of microgel solution was pipetted into Nunclon™ Sphera™ U-well 96-well plates (Thermofisher scientific™, 174925) with 10.000 cells in iPS-brew with 10 µM ROCK-inhibitor at a concentration of 10 µM for the first 24 hours. The volume was filled up to 150 µL per well, and the well plate was centrifuged for 4 min at 300 rcf using an Eppendorf Centrifuge 5810 R. After 24 hours media was changed to iPS-brew without inhibitor present. After 48-hours of iPSC-microgel scaffold formation, we started with the sensory or motor neuron differentiation, applying the same media protocol as described in the previous paragraph.

### Immunochemical and Molecular Staining

Cellular scaffolds were fixed with 4 % PFA for 45 minutes and washed with 1x PBS pH 7.4 three times, and afterward incubated in a solution containing 4 % BSA (SEQENS) and 0.1 % Triton X-100 (Sigma Aldrich) for 1 hour to permeabilize the cell membrane and block. Before the staining procedure, samples were washed three times with PBS. Primary staining of TUBB3 was used for all neuronal structures (Cell Signaling Technologies, anti-rabbit, 1:500 in 1% BSA and 0.1% Triton X-100 in PBS), SMI32 for motor neuron structures (Biolegend, anti-mouse, 1:500 in 1% BSA and 0.1% Triton X-100 in PBS), and Peripherin for sensory neuron structures (Santa Cruz Biotechnology, anti-mouse, 1:500 in 1% BSA and 0.1% Triton X-100 in PBS) over night at 4°C. Excess antibody is removed by washing the samples 3 times for 30 min with PBS, followed by incubation with the secondary antibodies (Alexa Fluor 568 goat anti-rabbit; Invitrogen, 1:200 or Alexa Fluor 647 goat anti-mouse; Invitrogen, 1:200) in PBS for 1 hour at RT. The samples are washed 3 times with PBS, and DAPI (1:200) in PBS is added for 10 min. After washing again thrice, the samples are stored in PBS, in the dark at 4°C until imaging. The same procedure is used for the other primaries: ChAT (Merck, anti-goat, 1:500), S100 (Invitrogen, anti-rabbit, 1:500), Glial Fibrillary Acid Protein (GFAP, Dako, anti-rabbit, 1:500), anti-O4 Antibody, clone 81 (Merck, anti-mouse, 1:500), and their corresponding secondaries: Alexa Fluor 647 goat anti-mouse (Invitrogen, 1:200), Alexa Fluor 568 goat anti-rabbit (Invitrogen, 1:200), and Alexa Fluor 488 donkey anti-goat (Invitrogen, 1:200).

### Sample clearing

For some samples, a tissue clearing was performed to allow better resolution and depth penetration for maximum projection recordings. No area or shape analysis was performed on cleared samples, since the clearing reagent can instigate size changes. Samples were cleared after immunofluorescence staining. Samples were incubated in an increasing concentration series of Isopropanol. The samples were incubated for 15 minutes, respectively, in 30 %, 50 %, and 70 % isopropanol solution (prepared with distilled water). Afterwards, the sample was incubated 2 times for 15 minutes in 100 % technical isopropanol solution. All Isopropanol steps were performed on ice. Before the removal of the last isopropanol step, the samples were warmed to room temperature (RT). The step of bringing the samples to RT was very important, since the later added ethyl cinnamate has a melting temperature of 6-8 °C, causing it to freeze if the samples are still on ice. The resulting ice crystals cause sample damage. After removal of the isopropanol, the samples were covered with pure ethyl cinnamate (ECi) and incubated for 20 minutes. Subsequently, samples could be directly imaged or stored in the ECi solution. Sample storage needs to be at RT to ensure the liquid state of the ECi.

### Image processing and Analysis

The samples are imaged using a Perkin Elmer Opera Phenix Plus with a 20x air or a 40x water objective. These images are captured using the appropriate excitation wavelengths, and the emission signals are captured using detection sCMOS sensor cameras.

Image analysis was conducted with the ImageJ software. Area analysis of scaffolds was performed, generating a 16-bit image, converting this image into a binary image using the default threshold method, and subsequently measuring the white pixel area. The white pixel area was transformed into spheroid area through scale definition according to the scale bar of the microcopy images. Roundness of the samples was also measured with ImageJ software using the same method as described for the area. Roundness was defined as:

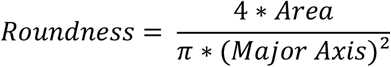

To enable easier analysis, macros were implemented for automated area and roundness analysis locally in the ImageJ software.

### Statistical analysis

Statistical analysis and graphs were produced using GraphPad Prism 9 software. Box plots represent boxes and whiskers from minimal to maximal values, including all data points. The performed statistical analysis for each data set is reported in the respective figure caption. For statistical analysis, the P-values were represented in the figures.

## Acknowledgments

The authors gratefully acknowledge funding from the Canadian government via the New Frontiers in Research Fund (NFRF), NFRFT-2020-00238, within the project Mend the Gap, the DFG weave (LA 3606/9-1 AOBJ: 719038), AnisoMatic IGF 01IF23480N, the European Union’s Horizon Europe research and innovation program under the grant agreement number 101191649 (NEOLIVER), and the European Research Council within the ERC-2021-COG project 101043656, Heartbeat.

This work was supported by the Core Facility “Two-Photon Imaging” and the IZKF High Content Screening Facility, the Core Facilities of the Interdisciplinary Center for Clinical Research (IZKF) Aachen within the Faculty of Medicine at RWTH Aachen University.

## Author’s contributions

L. K., C. B., J. L. G. N., and L. D. L. conceived the initial idea. L. K., C. B., J. L. G. N., and L. D. L. designed all experiments, which L. K. and C. B. performed. L. K. and C. B. imaged the experiments. M. M. and G. R. produced the microgels. L. K. and C. B. wrote the manuscript, which J. L. G. N. and L. D. L. corrected. All authors read and provided feedback on the manuscript.

## Data Availability Statement

The data that support the findings of this study are available from the corresponding author upon reasonable request.

