## Supplementary information for "Differentiation and maturation of iPSC-derived motor and sensory neurospheres using biomodified PEG-based microgels"

### Supporting Information

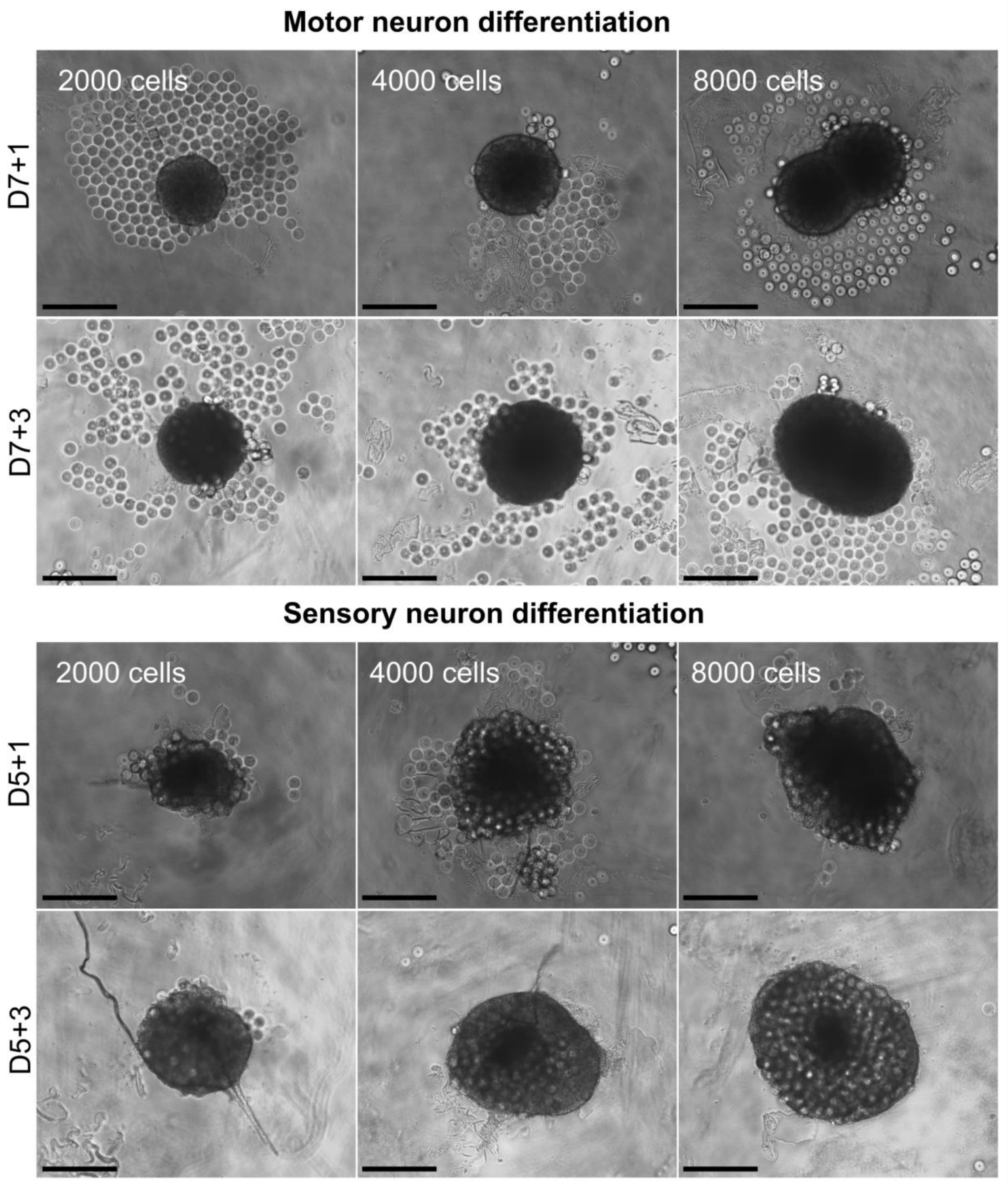


**Figure SI1:** Initial experiment showing precursor motor and sensory neuron spheres of different sizes added after 7 days (motor) and 5 days (sensory) to VTN-coated microgels. The first row is after 1 day of combined cultivation, and the second row is after 3 days. The scale bar is 500 µm.

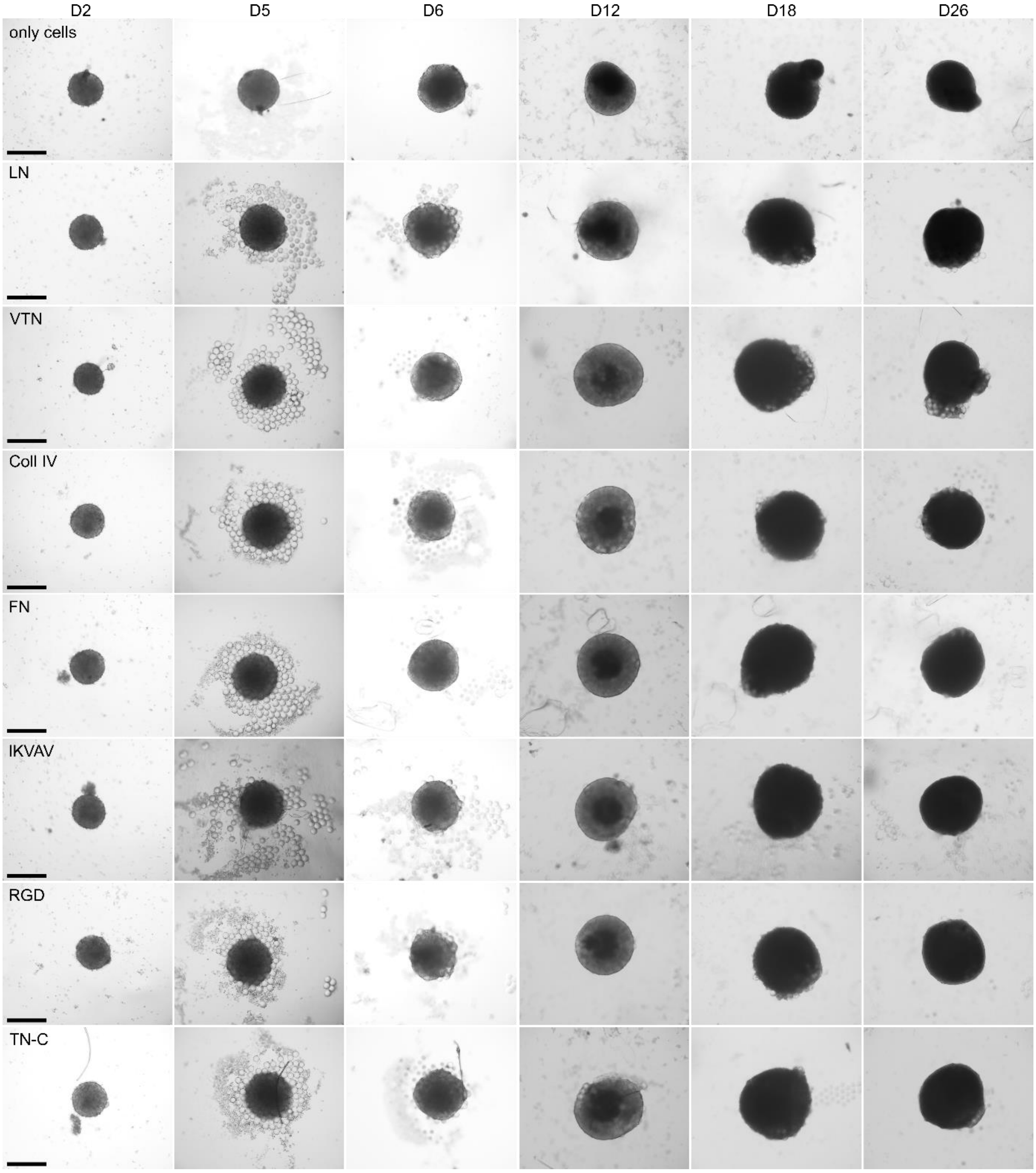


**Figure SI2:** Sensory neurospheres with different biopeptide-coated microgels over 26 days of cultivation time. Brightfield images on day 2, 5, 6, 12, 18, 26 of the interaction of neural crest spheroids with different biopeptide-coated microgels (laminin, VTN, CollIV, FN, IKVAV, RGD, Tenascin C) and only-cells during their differentiation and maturation to sensory neurospheres over 26 days. The scale bar is 500 µm.

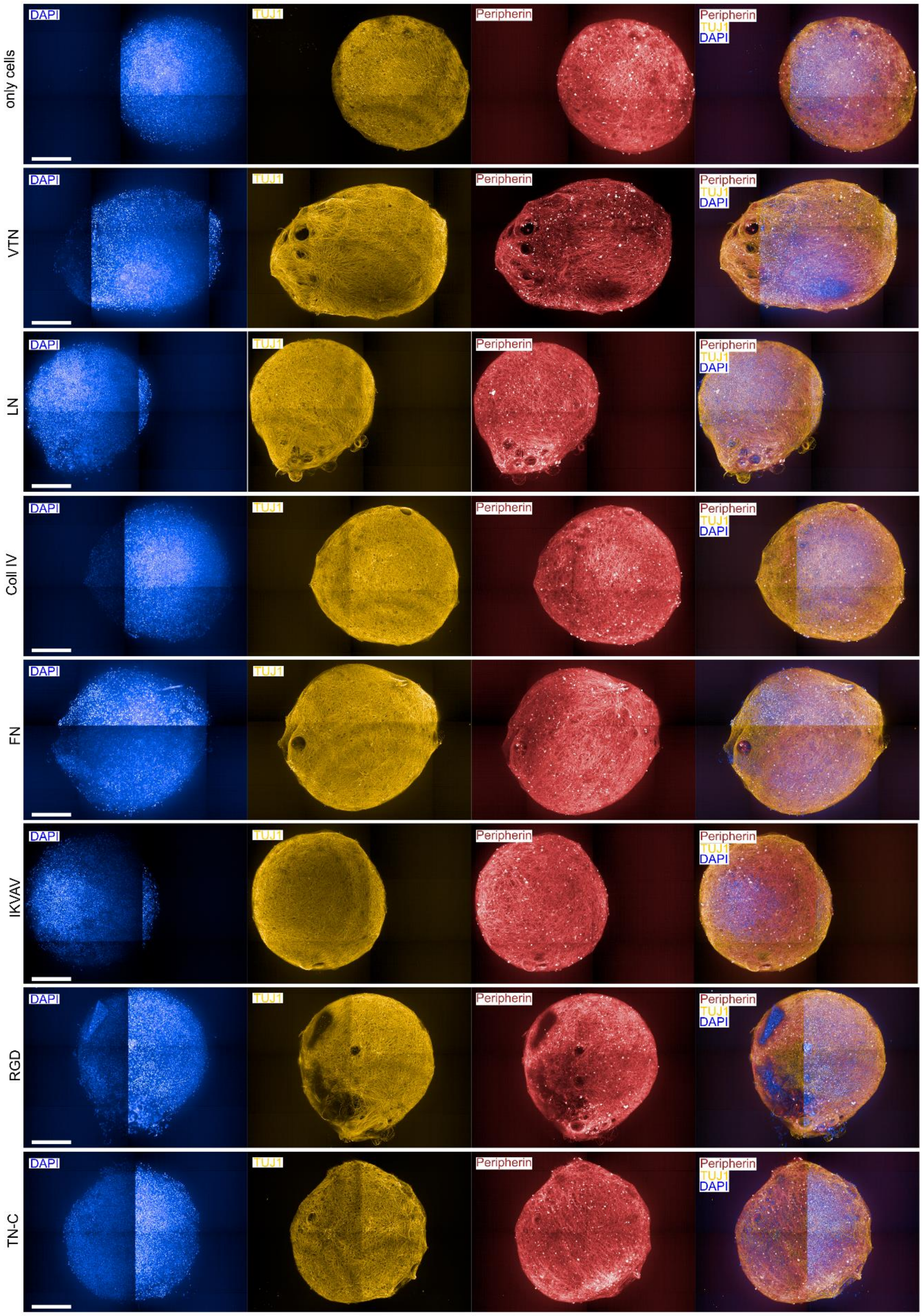


**Figure SI3:** Sensory neurospheres with different biopeptide-coated microgels after 26 days of cultivation time. Immunofluorescent images of DAPI (blue), TUJ1 (yellow), peripherin (red), and their corresponding merged image of the interaction of sensory neurospheres with different biopeptide-coated microgels (laminin, VTN, CollIV, FN, IKVAV, RGD, Tenascin C) after 26 days of culture. The scale bar is 200 µm.

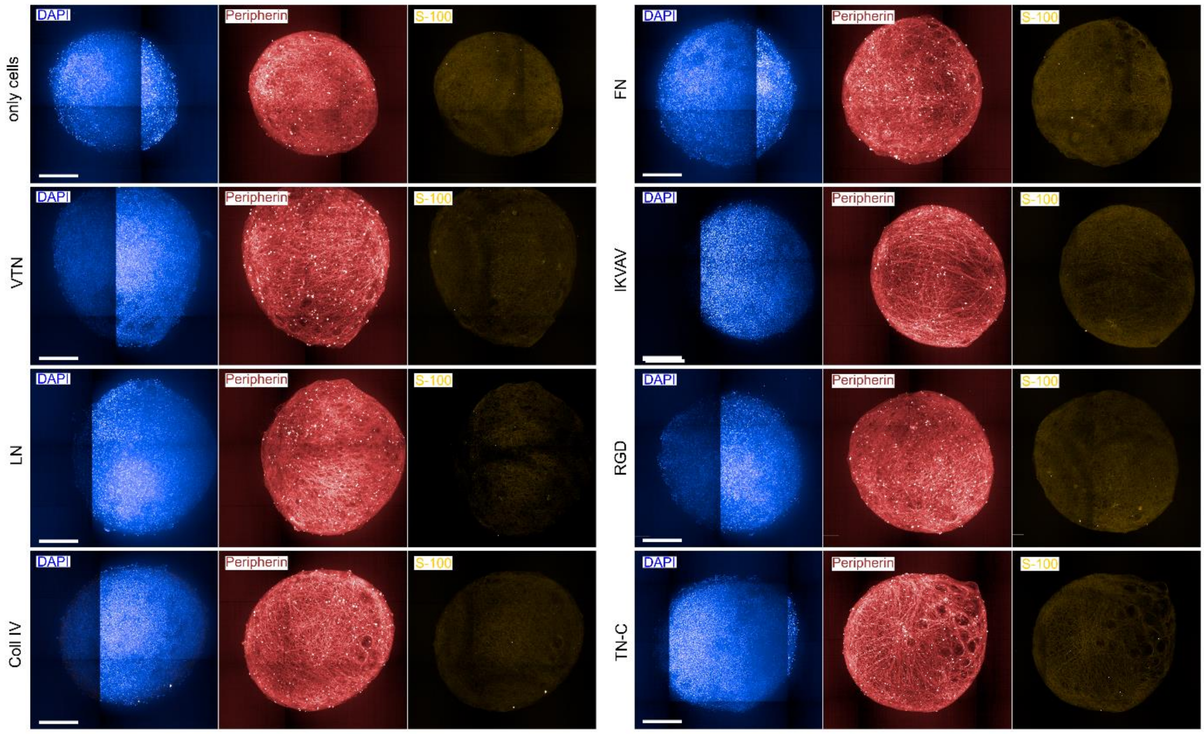


**Figure SI4:** Sensory neurospheres with different biopeptide-coated microgels after 26 days of cultivation time. Immunofluorescent images of DAPI (blue), peripherin (red), and S100 (yellow) of the interaction of sensory neurospheres with different biopeptide-coated microgels (LN, VTN, CollIV, FN, IKVAV, RGD, TN-C) and only-cells after 26 days of culture. The scale bar is 200 µm.

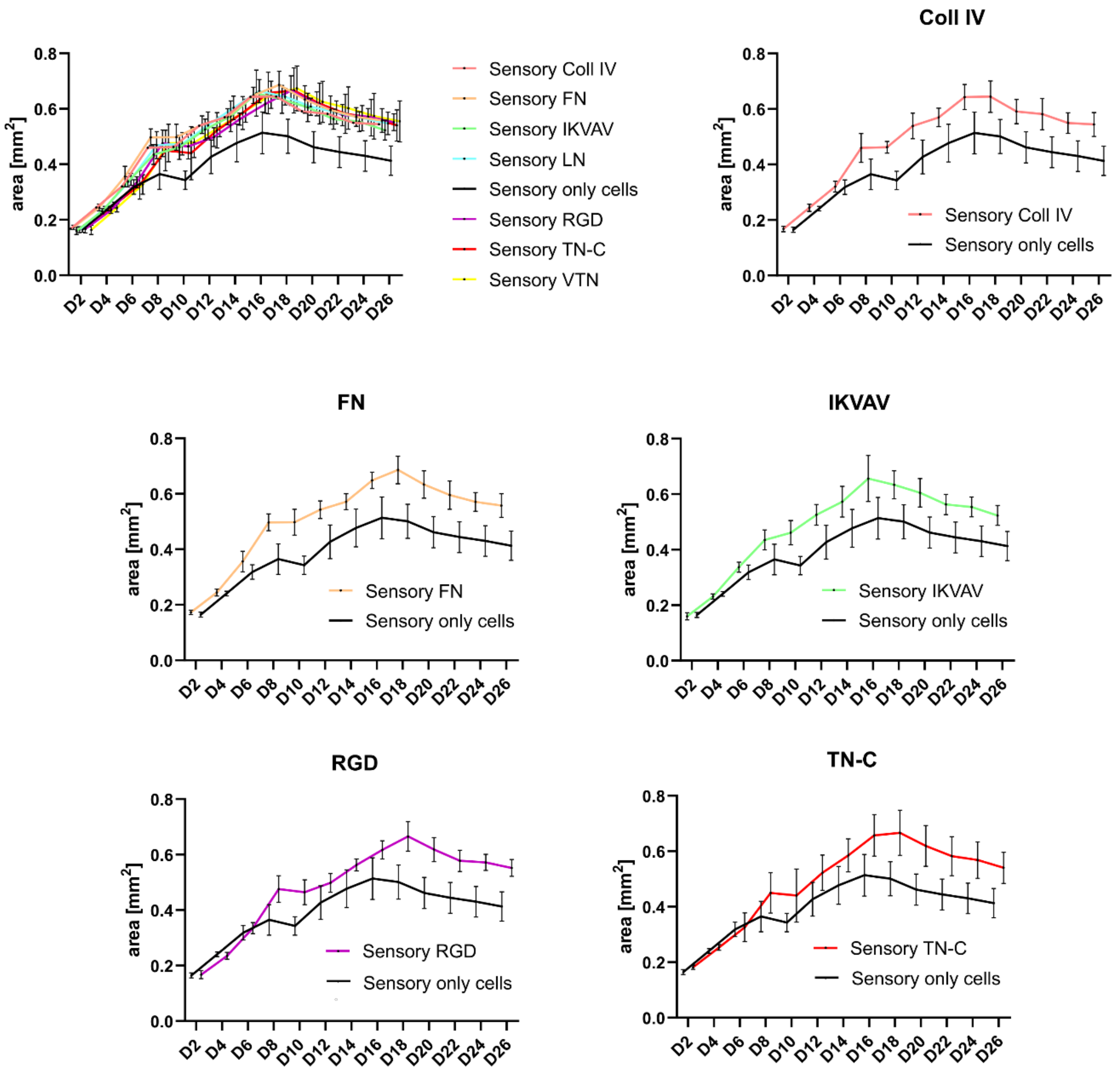


**Figure SI5:** Area analysis of the sensory neurospheres over 26 culturing days, comparing spheroids cultured with CollIV-(light red), FN-(orange), IKVAV-(cyan), with LN-(green), RGD (violet), TN-C-(red), VTN-(yellow) coated microgels, and only-cells (black).

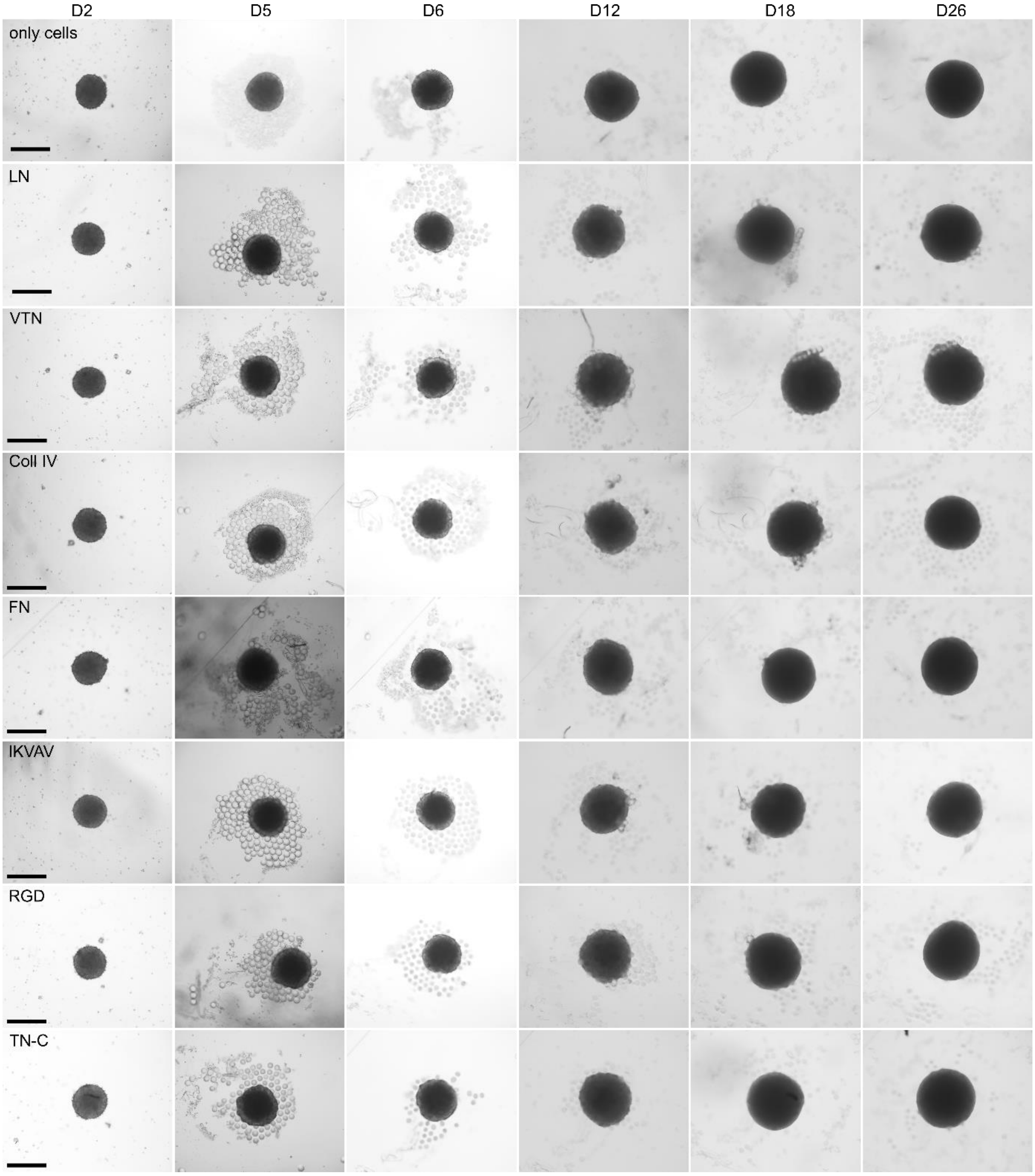


**Figure SI6:** Motor neurospheres with different biopeptide-coated microgels over 26 days of cultivation time. Brightfield images on day 2, 5, 6, 12, 18, 26 of the interaction of motor precursor spheroids with different biopeptide-coated microgels (LN, VTN, CollIV, FN, IKVAV, RGD, TN-C) and only-cells during their differentiation and maturation to motor neurospheres over 26 days. The scale bar is 500 µm.

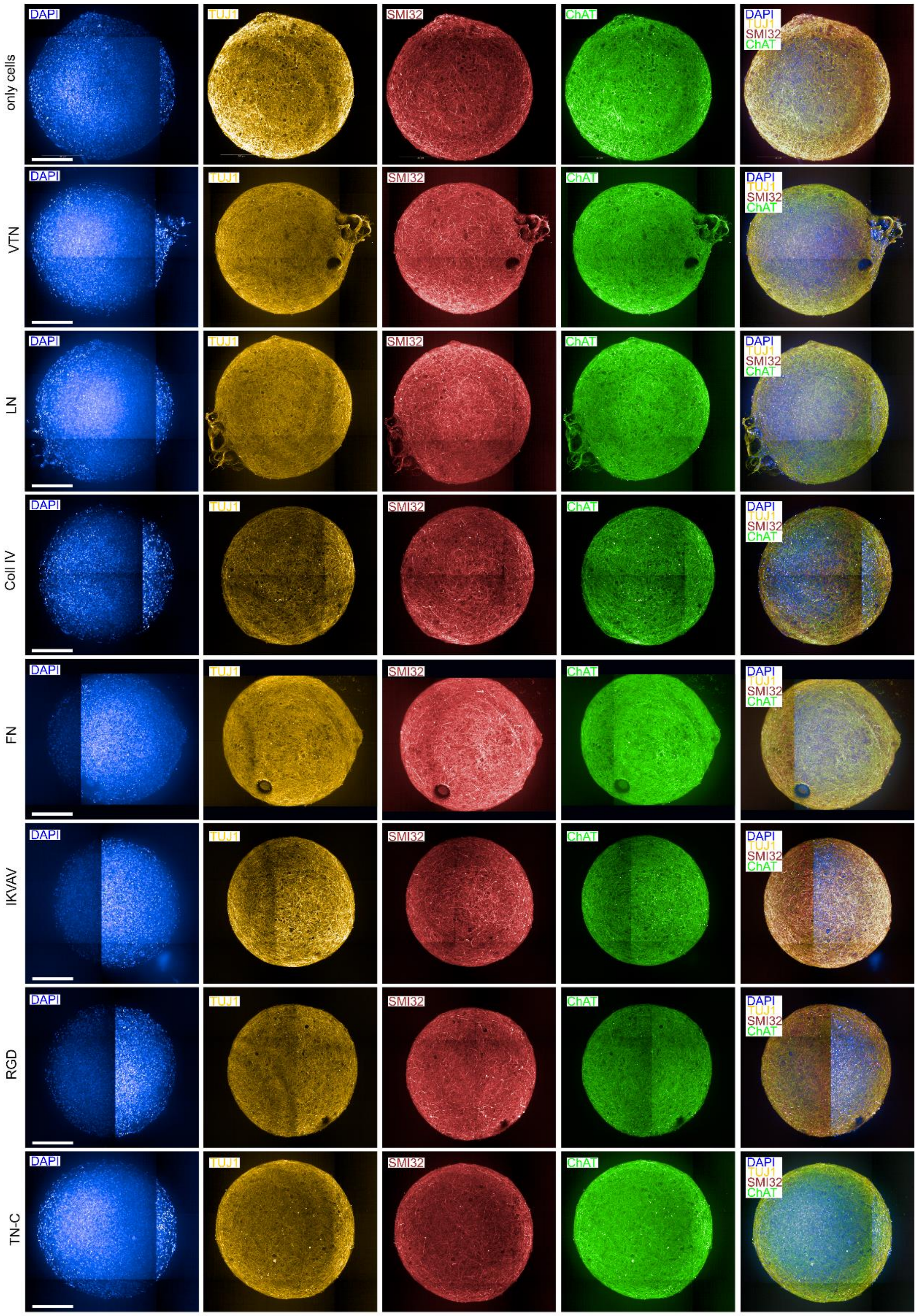


**Figure SI7:** Motor neurospheres with different biopeptide-coated microgels after 26 days of cultivation time. Immunofluorescent images of DAPI (blue), TUJ1 (yellow), SMI32 (red), ChAT (green), and their corresponding merged image of the interaction of motor neurospheres with different biopeptide-coated microgels (LN, VTN, CollIV, FN, IKVAV, RGD, TN-C) and only-cells after 26 days of culture. The scale bar is 200 µm.

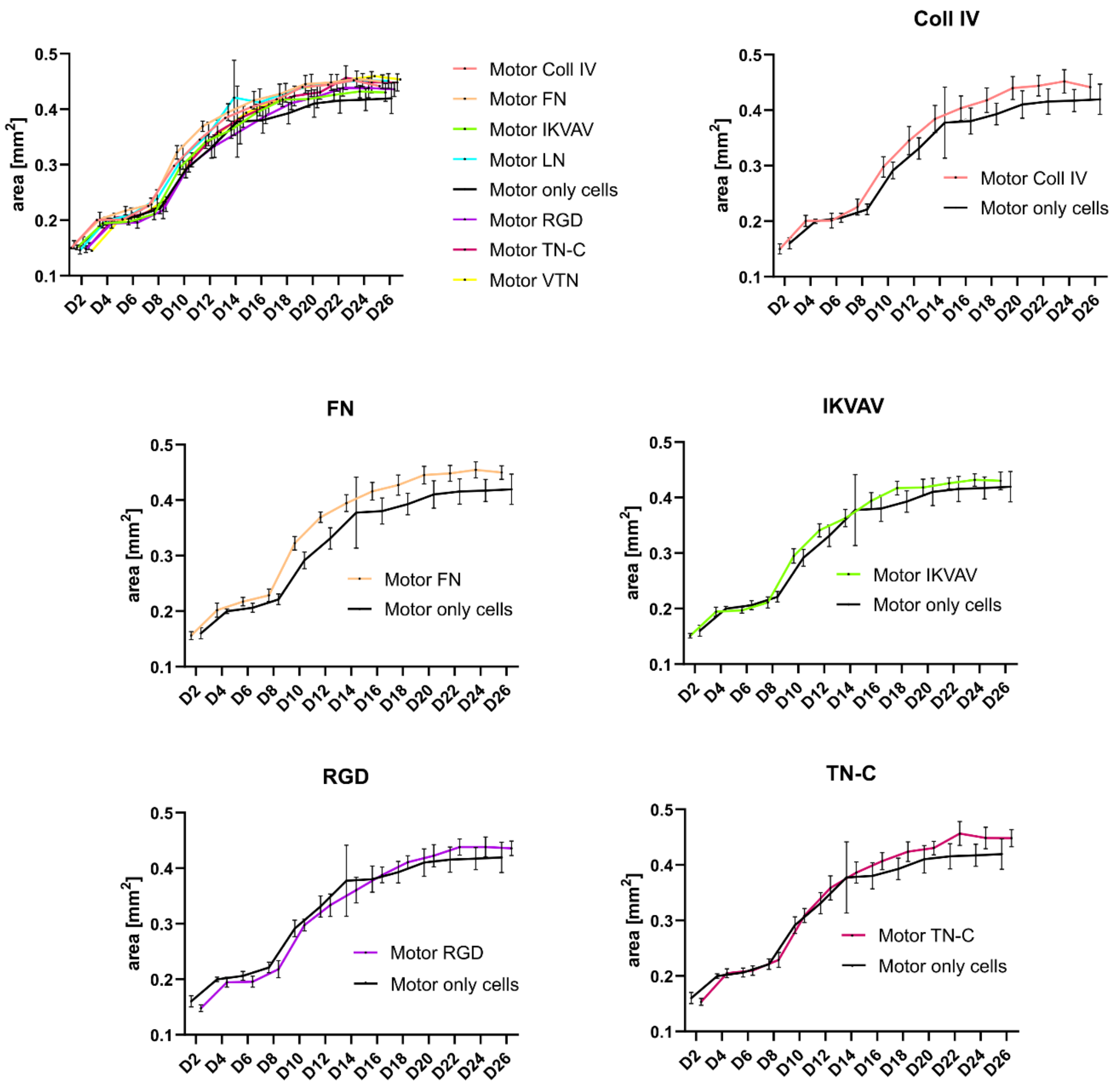


**Figure SI8:** Area analysis of the motor neurospheres over 26 culturing days, comparing spheroids cultured with CollIV-(light red), FN-(orange), IKVAV-(cyan), with LN-(green), RGD (violet), TN-C-(red), VTN-(yellow) coated microgels, and only-cells (black).

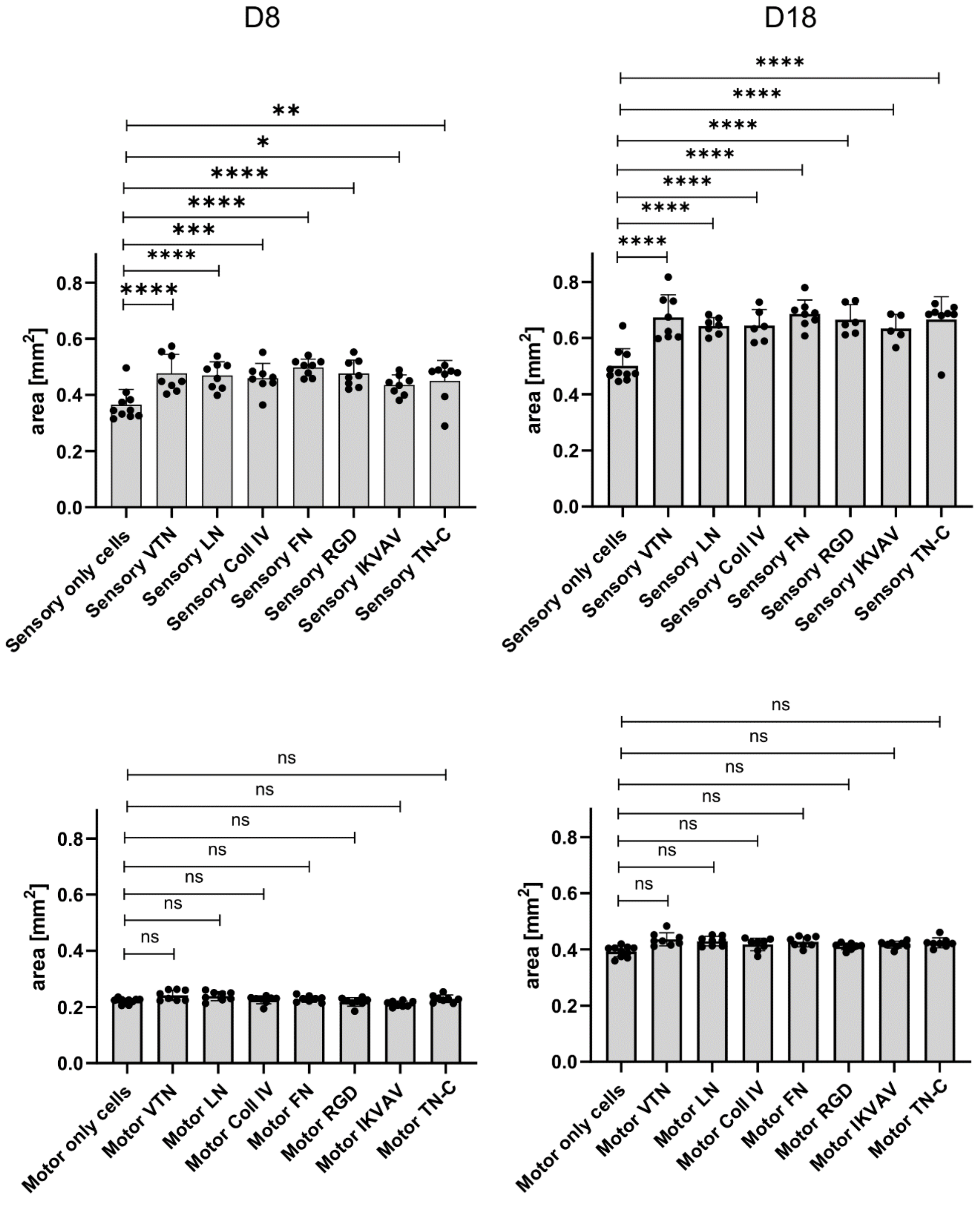


**Figure SI9:** Area comparison of the motor and sensory neurospheres at day 8 and day 18, comparing spheroids cultured with CollIV-, FN-, IKVAV-, with LN-, RGD, TN-C, VTN-coated microgels, and only-cells.

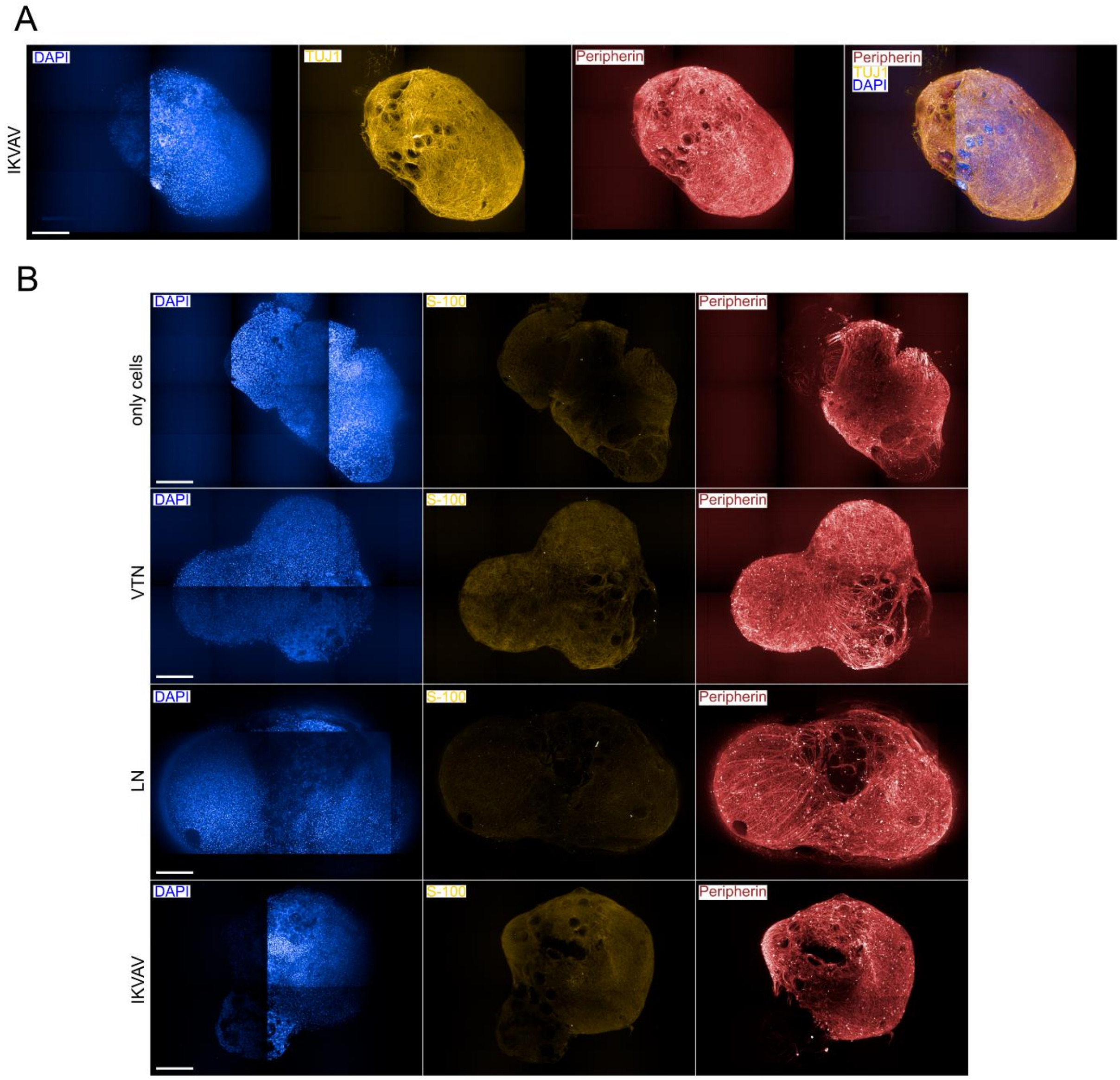


**Figure SI10:** Sensory neurospheres from iPSCs with different biopeptide-coated microgels after 26 days of cultivation time. A, B. Immunofluorescent images of DAPI (blue), peripherin (red), and TUJ1(A, yellow) or S100 (B, yellow) of the interaction of sensory neurospheres with different biopeptide-coated microgels (LN, VTN, IKVAV) and only-cells after 26 days of culture. The scale bar is 200 µm.

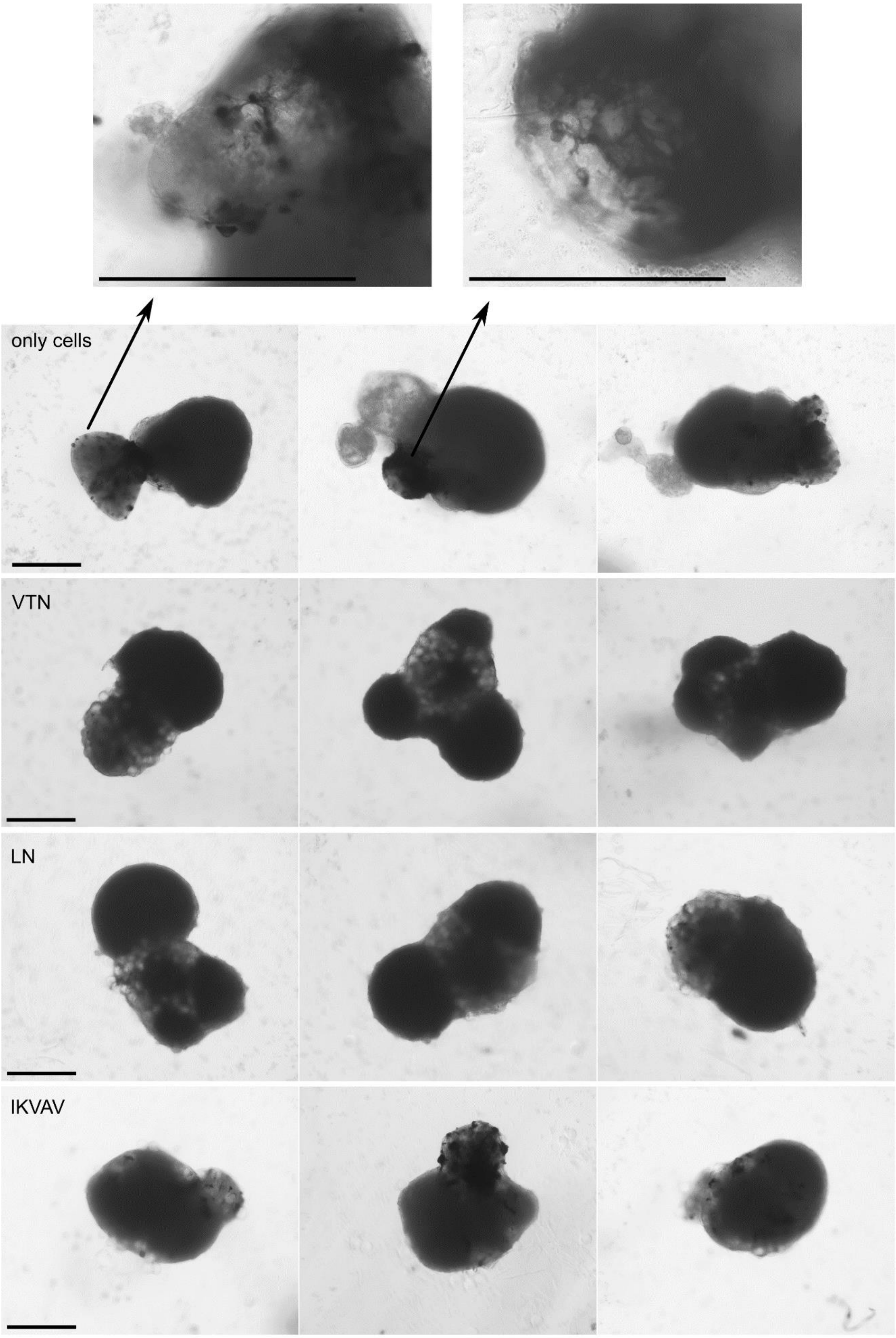


**Figure SI11:** Brightfield images of sensory neurospheres with different biopeptide-coated (VTN, LN, IKVAV) microgels and only-cells after 26 days of cultivation time, indicating the presence of melanocytes in the spheroids. The scale bar is 500 µm.

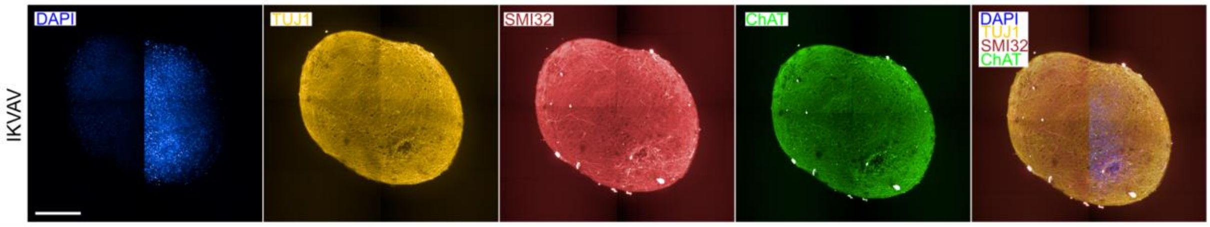


**Figure SI12:** Immunofluorescent images of DAPI (blue), TUJ1 (yellow), SMI32 (red), ChAT (green), and their corresponding merged image of the interaction of motor neurospheres with IKVAV-coated microgels after 26 days of culture. The scale bar is 200 µm.

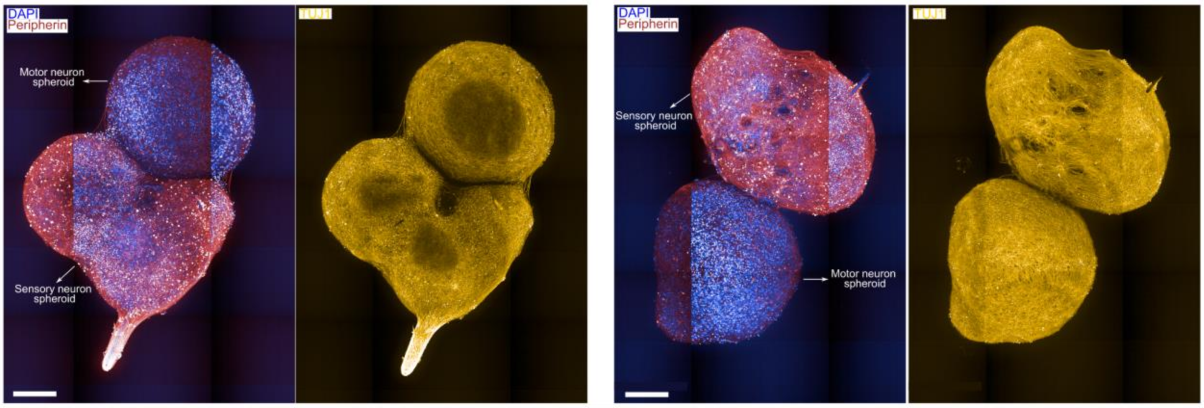


**Figure SI13:** Immunofluorescent images of DAPI (blue), TUJ1 (yellow), and peripherin (red) of the interaction of motor neurospheres with sensory neurospheres after 26 days of culture. The scale bar is 200 µm.
